# Diversity in early animal communities of the Ediacaran, Avalon assemblage

**DOI:** 10.64898/2026.10.28.762103

**Authors:** Shujie Chang, Nile P. Stephenson, Andrea Manica, Emily G. Mitchell

## Abstract

Diversity is an emergent property of macroecological and macroevolutionary processes. The Avalon assemblage preserves near-census snapshots of Ediacaran communities, enabling tests of the drivers of diversity. However, it remains unclear how alpha diversity varied among surfaces or whether variation reflected uneven sampling, abiotic context or biotic processes. We analysed diversity for 11,342 specimens across 22 surfaces and examined abiotic and biotic correlates. Despite compositional variation, most surfaces fell into four community types dominated by *Fractofusus andersoni*, *F. misrai* and/or *Pectinifrons*, *Bradgatia*, or upright fronds, with potential substructure within the upright-frond type, whereas Clapham’s Pigeon Cove and Shingle Head communities, from the oldest and youngest formations, respectively, may indicate greater community differentiation near the sampled stratigraphic limits. Sampling effort varied widely, yet taxon–area curves showed richness was well captured on 12 of 22 surfaces, and Shannon and Simpson diversity remained consistent across sampling completeness and taxonomic treatments. Diversity showed no clear associations with locality or stratigraphic affiliation, but differed among community types and declined with increasing vertical segregation. Overall, these results suggest that, among the predominantly Newfoundland communities sampled, Avalon alpha diversity may have been shaped more by local community assembly and organisation than by broader geographic, environmental and temporal context.

## Introduction

Diversity is an emergent property of macroecological and macroevolutionary processes operating across multiple spatial and temporal scales. Across space, environmental heterogeneity and differences in dispersal or connectivity among communities can shape local diversity by influencing which taxa can reach, establish and coexist under local conditions^1–3^. Through time, changes in regional species pools associated with speciation and extinction can alter the taxa available to local communities and, consequently, their diversity^4,5^. At local scales, biotic processes including recruitment, competition, facilitation and niche differentiation can further shape diversity by influencing which taxa coexist and how resources are partitioned^6–9^. Although these processes are well studied in modern ecological systems, their relative roles in shaping diversity in early animal communities remain poorly understood.

The emergence of animal communities during the Ediacaran marked a major transition in the history of life, providing a window into the ecological and evolutionary context of early animal diversification^10–13^. The Avalon assemblage (∼574–560 Ma) preserves deep-marine seafloor communities in situ as near-census snapshots on fossiliferous surfaces in Newfoundland, Canada, and Charnwood Forest, UK^12,14,15^. This exceptional preservation provides a rare basis for comparing diversity among early animal communities^12,16,17^. Across these fossiliferous surfaces, community composition varies markedly, as reflected in high between-surface beta diversity^17,18^.

Previous research on Avalonian communities has revealed patterns across multiple scales, from metacommunity organisation to among-surface differences in composition and local interactions. At the metacommunity scale, Avalonian communities show relatively limited ecological specialisation and taxon segregation^19^. Among communities, recurring assemblage types emerge from differences in composition and span different successional stages^16,17,20^. Community structure also varies in the extent of vertical partitioning of resource-acquisition space within the water column, a pattern known as tiering^17,21,22^. Local spatial patterns are often consistent with neutral processes, potentially in part because competitive interactions can be weakened by reproductive mode^23,24^. Despite this increasingly detailed understanding of Avalonian community organisation, how alpha diversity varies among communities, and the extent to which that variation is associated with broader spatial, temporal and environmental context or with local community organisation, remains unresolved.

For Avalonian communities, these dimensions can be examined using a set of observable correlates. Stratigraphy provides a broad proxy for temporal and environmental context^15,25^, while locality captures geographic context and may also integrate differences in environmental conditions or connectivity among communities^2^. At the community level, composition and tiering represent complementary dimensions of biotic organisation: composition summarises recurring differences in taxonomic identity, relative abundance and dominance, whereas tiering describes the vertical partitioning of resource-acquisition space and thus one dimension of niche differentiation^17,21,22,26^. Together, these correlates provide a framework for evaluating variation in alpha diversity across broader context and local community organisation.

Before these relationships can be evaluated, it is necessary to establish whether observed differences in diversity reflect underlying differences among communities rather than variation in sampling effort^27,28^. Avalonian fossiliferous surfaces differ substantially in both mapped area and the number of recorded specimens^17^, potentially affecting the recovery of taxa and therefore observed diversity^29^. Taxon–area accumulation curves provide a means of assessing whether the accumulation of taxonomic richness approaches an asymptote with increasing sampled area^28,30^, while extrapolated diversity estimates approximate the diversity expected under more complete sampling^27^. Together, these approaches provide a basis for distinguishing signals due to sampling effects from broader ecological patterns of diversity variation.

Here, we use 22 Avalonian communities to systematically assess variation in alpha diversity. We first evaluate sampling completeness and quantify the extent to which variation in specimen number and mapped area is associated with observed diversity. We then compare observed and extrapolated diversity estimates before testing whether diversity covaries with broader spatial, temporal and environmental context, represented by stratigraphy and locality, and with community-level biotic organisation, represented by community composition and tiering, thereby identifying which dimensions of ecological context covary most consistently with Avalonian diversity.

## Methods

### Study data

We compiled data from 22 Avalonian fossiliferous surfaces. These comprised Brasier, Bristy Cove, Clapham’s Pigeon Cove, D Surface, the E and G surfaces at both Mistaken Point and Watern Cove West (E) and East (G), Lower Mistaken Point, Pizzeria, Shingle Head and St. Shotts from the Mistaken Point Ecological Reserve; Bishop’s Cove, Green Point and Upper Island Cove from the Spaniard’s Bay area; The Factory (H31), Goldmine, H5, H14 Johnson, H26 Primocandelabrum and Caplin Gulch from the Discovery Geopark area, all in Newfoundland, Canada; and Bed B from Charnwood Forest, Leicestershire, UK (Table S1). Within Newfoundland, the studied surfaces span the Drook, Briscal, Mistaken Point, Trepassey and Fermeuse formations within the Mistaken Point Ecological Reserve; the Mistaken Point and Trepassey formations in the Spaniard’s Bay area; and the Murphy’s Cove, Catalina and Port Union members and the Fermeuse Formation of the Discovery Geopark succession^31–34^; Bed B is from the Bradgate Formation in Charnwood Forest, Leicestershire, UK (Table S1). The surface data comprised published datasets^17,35^.

Surface mapping used a combination of photogrammetry, light detection and ranging (LiDAR) and, where surface geometry and fossil preservation permitted, high-resolution laser-line probe (LLP) scanning^23^. LiDAR scans generally produced surface models with a mean resolution of up to 0.5 mm, while LLP scanning provided higher-resolution morphological data. In particular, Mistaken Point G Surface was scanned using a FARO ScanArm LLP v6 at a resolution of 40 μm. The steeply dipping E and G surfaces at Watern Cove could not be scanned using the LLP and were instead scanned using a FARO 330 LiDAR scanner at 1 mm resolution, supplemented by photogrammetry. Photogrammetric maps were used for H14 Johnson, Bristy Cove and St. Shotts, where fossils could be distinguished from colour contrasts within the surfaces, while Bed B was digitised from reflectance transformation images of casts^36^. Three-dimensional surface models and orthomosaics were generated from the photogrammetric data in Agisoft Metashape v1.7.5, and fossils were digitised in Inkscape v1.3 to produce two-dimensional vector maps. The resulting specimen-level datasets contained taxonomic identifications, spatial coordinates, morphological measurements (including frond length and stem length).

To account for tectonic deformation affecting many Avalonian fossiliferous surfaces^14,15^, surface maps were retrodeformed before analysis to restore specimen morphologies and spatial positions. Deformation was quantified from the systematic elongation of holdfast discs, which were assumed to have been approximately circular before tectonic compression^15^. For each surface, the retrodeformation factor was calculated as the mean ratio of the long to short axes of suitable frondose holdfast discs larger than 2 cm since smaller discs can carry sufficient measurement errors to create noise. Surface maps were then rescaled along the inferred direction of shortening, as indicated by the short axes of the deformed discs. No retrodeformation was applied where suitable discs were unavailable or where the estimated factor was less than 1.05. Retrodeformation factors and resulting mapped areas are reported in Table S2.

Taxonomic identifications were reviewed and standardised across surfaces before analysis. The dataset included both formally described taxa and informally named morphotypes that could be consistently distinguished across surfaces. We retained specimens that could be reliably assigned to one of the taxonomic units used in the analyses and excluded organ taxa, taphomorphs and effaced fronds, the latter representing eroded frondose specimens that could not be identified to genus level. Because species-level boundaries remain unresolved for some taxa and diagnostic characters were not consistently preserved across surfaces, the main dataset was standardised to an approximately genus-level taxonomic resolution. Specimens of *Charniodiscus* and *Primocandelabrum* were pooled at genus level, while *Aninoides coombsorum*^37^ was grouped with *Beothukis* within the broader “beothukiid” taxonomic group. *Bradgatia* was retained at genus level in both taxonomic treatments because species-level differentiation within the genus remains unresolved, particularly for the Newfoundland material^38^ (see also Extended Data Table 3 in Ref. 17). The final analytical dataset covered a total mapped area of 853.45 m² and comprised 11,342 specimens assigned to 27 taxonomic units (Table S2).

### Community-type classification

To assess variation in community composition among surfaces, pairwise Bray–Curtis dissimilarities were calculated from taxon relative abundances using the vegan package^39^. Compositional relationships among surfaces were visualised using two-dimensional non-metric multidimensional scaling (NMDS), with 200 random starts. Average-linkage hierarchical clustering (UPGMA) was then applied to the same Bray–Curtis dissimilarity matrix. Cluster stability was assessed using multiscale bootstrap resampling with 1,000 bootstrap replicates in the pvclust package (v2.2.0)^40^, with approximately unbiased (AU) *p*-values used to quantify support for each internal node.

Candidate numbers of clusters, ranging from *k* = 2 to one fewer than the number of surfaces, were evaluated by cutting the dendrogram at each value of *k* and calculating the corresponding mean silhouette width^41^ from the Bray–Curtis dissimilarity matrix using the cluster package (v2.1.8.2)^42^. The clustering and ordination results were interpreted alongside dominant-taxon composition and the four previously described community types^17^: *Bradgatia*-dominated, *Fractofusus andersoni*-dominated, *F. misrai* and/or *Pectinifrons*-dominated, and frond-dominated communities.

### Tiering

To quantify community-level tiering, we reconstructed the in-life vertical extent of each specimen above the seafloor. Because the relationship between preserved dimensions and in-life height depends on growth habit, each taxonomic unit was classified as upright or reclining. For upright taxa, height was calculated as frond length plus stem length when a stem was present. For taxa classified as reclining, vertical extent above the seafloor was estimated as one-third of frond width ^22^. Fronds were treated as the primary resource-acquisition structures, whereas stems were interpreted as providing mainly structural support^43–45^. Therefore, stem length was used to position the frond above the seafloor but was excluded from the zone by which the animal would capture food and other nutrients, namely the uptake zone. Community-level tiering was quantified using the uptake-zone formulation of Distinct Vertical Stratification (DVS), with taxon-specific values weighted by each taxon’s relative abundance within the surface^17,22^. DVS ranges from 0, representing complete overlap in vertical resource-acquisition space, to 1, representing perfect tiering. To assess whether community-level tiering differed among the community types, DVS values were compared using a Kruskal–Wallis test.

### Sampling completeness

To assess whether the mapped extent of each surface was sufficient to capture most of its taxonomic richness, we generated sample-based taxon–area accumulation curves^28,30^. Each mapped surface was divided into a 10 × 10 grid, and specimens were assigned to the quadrats in which they occurred using the spatstat package (v3.6.1)^46,47^ in R (v4.6.0)^48^. Taxonomic richness was accumulated across quadrats in random order using the specaccum function with 10,000 permutations in the vegan package (v2.7.5)^39^. For each accumulation step from one to 100 quadrats, we recorded the mean taxonomic richness and its standard deviation, and calculated cumulative sampled area as the number of accumulated quadrats multiplied by the mean quadrat area for that surface. We then fitted four nonlinear asymptotic models to mean taxonomic richness as a function of sampled area: two- and three-parameter Michaelis–Menten models (MM.2 and MM.3) and two- and three-parameter asymptotic regression models (AR.2 and AR.3), implemented using the drc package (v3.0.1)^49^. The model with the lowest Akaike Information Criterion (AIC) was selected for each surface, and its upper asymptote was taken as the fitted asymptotic taxonomic richness. A surface was classified as saturated when its observed richness reached at least 80% of the fitted asymptotic richness.

### Diversity estimation

For each surface, we calculated four observed diversity metrics from taxon abundances: taxonomic richness, Shannon entropy, the Gini–Simpson index and Pielou’s evenness. To estimate diversity under more complete sampling, we applied abundance-based asymptotic estimators for taxonomic richness, Shannon entropy and Gini–Simpson index using the iNEXT package (v3.0.2)^27,50^. To obtain a corresponding asymptotic estimate of Pielou’s evenness for comparison with the observed value, we calculated it as

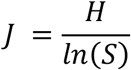

where *H* is asymptotic Shannon entropy and *S* is asymptotic taxonomic richness. Standard errors for *J* were approximated from the standard errors of *H* and *S* using the delta method^51^. These standard errors were then used to calculate 95% confidence intervals under a normal approximation, with intervals truncated to the range 0–1.

### Sampling effects

To test whether cross-surface diversity patterns were associated with sampling effort, we fitted separate linear regressions of diversity against specimen count and retrodeformed area. For each sampling measure, regressions were fitted using observed diversity among surfaces classified as saturated, observed diversity across all surfaces, and estimated diversity across all surfaces. These analyses respectively evaluated patterns in a subset with stronger evidence of sampling completeness, in the full observed dataset, and after asymptotic estimation of unobserved diversity. We additionally used Wilcoxon rank-sum tests to compare retrodeformed area and specimen count between saturated and non-saturated surfaces.

### Candidate correlates of diversity

We tested locality, stratigraphic unit, community type and DVS as candidate correlates of diversity. Candidate-correlate analyses likewise used observed diversity among saturated surfaces, and observed and estimated diversity across all surfaces. Stratigraphy was represented using two categorical classification schemes. The local-unit scheme used the region-specific stratigraphic assignments, whereas the correlated scheme grouped these into their formation-level equivalents based on regional lithostratigraphic correlations (Table S1). Specifically, the Murphy’s Cove Member was grouped with the Mistaken Point Formation, and the Catalina and Port Union members with the Trepassey Formation^31,33^.

Locality, stratigraphic affiliation and community type were treated as categorical variables. To ensure a minimum level of replication, only categories represented by at least three surfaces were included in formal statistical comparisons. Comparisons between two groups were performed using Wilcoxon rank-sum tests, whereas comparisons among three or more groups were performed using Kruskal–Wallis tests. DVS was treated as a continuous variable and tested using linear regression. To evaluate the joint and potentially interactive effects of community type and DVS, we additionally fitted a global model containing DVS, community type and their interactions.

Candidate models comprising subsets of the global model were generated using the dredge function in the MuMIn package (v1.48.19)^52^. Candidate models were ranked by the small-sample-corrected Akaike information criterion (AICc) and compared using ΔAICc.

### Sensitivity analyses

In our dataset, the E and G surfaces were each represented by outcrops at Mistaken Point and Watern Cove West (E) and East (G) and were therefore treated as four analytical surfaces. Although the paired outcrops are separated by 0.68 km for the E Surface and 0.78 km for the G Surface, they represent repeated sampling of the same stratigraphic horizons and so do not constitute fully independent ecological replicates^35^. To assess whether their repeated representation influenced the results, we repeated the community-type classification and diversity analyses after excluding the smaller E Surface at Watern Cove West and G Surface at Mistaken Point datasets.

To assess sensitivity to taxonomic resolution, all relevant analyses were also repeated using the finest available taxonomic identifications within the “beothukiid”, *Charniodiscus* and *Primocandelabrum* groupings. This finer taxonomic resolution produced a finer but mixed-resolution treatment in which species-level assignments were retained where diagnostic morphology and preservation permitted, whereas specimens that could not be assigned confidently to species, including forms that may represent undescribed species, were retained at genus level. Because named species and potentially overlapping unresolved categories were treated as separate taxonomic units, this treatment could alter both the number and relative abundances of recognised taxa within surfaces and the shared taxon pool used in cross-surface community analyses. It was therefore used as a sensitivity analysis rather than as the primary taxonomic treatment.

To test the influence of these different taxonomic groupings, agreement between diversity metrics obtained under the two taxonomic treatments was quantified across surfaces using Lin’s concordance correlation coefficient (CCC)^53^, with 95% confidence intervals calculated using a z-transformation in the DescTools package (v0.99.60)^54^. CCC combines precision and accuracy by quantifying how closely paired estimates follow the 1:1 line of perfect concordance. CCC values range from −1 to 1, with values closer to 1 indicating stronger concordance. CCC values were interpreted using the following empirical categories: poor (<0.90), moderate (0.90–<0.95), substantial (0.95–<0.99) and almost perfect (≥0.99) agreement^55^.

## Results

### Community composition

Among the 22 communities, community composition varied markedly in both relative-abundance structure and the identity of dominant taxa. Taxonomic richness ranged from 2 to 18 taxa, with a mean of 7.4 taxa per community (SD = 4.0; Fig. 1a; Table S2). The most abundant taxon accounted for 31.5–99.5% of specimens within individual communities, ranging from relatively mixed to near-monodominant assemblages. When abundances were pooled across all communities, the rank–abundance distribution was strongly uneven, with *Fractofusus andersoni* and *F. misrai* substantially more abundant than most other taxa, followed by a long tail of low-abundance taxa (Fig. 1b). The same broad compositional patterns were retained under the finer-resolution taxonomic treatment, in which specimens within the “beothukiid”, *Charniodiscus* and *Primocandelabrum* groupings were resolved to species level where possible (Fig. S1).

**Figure 1.**
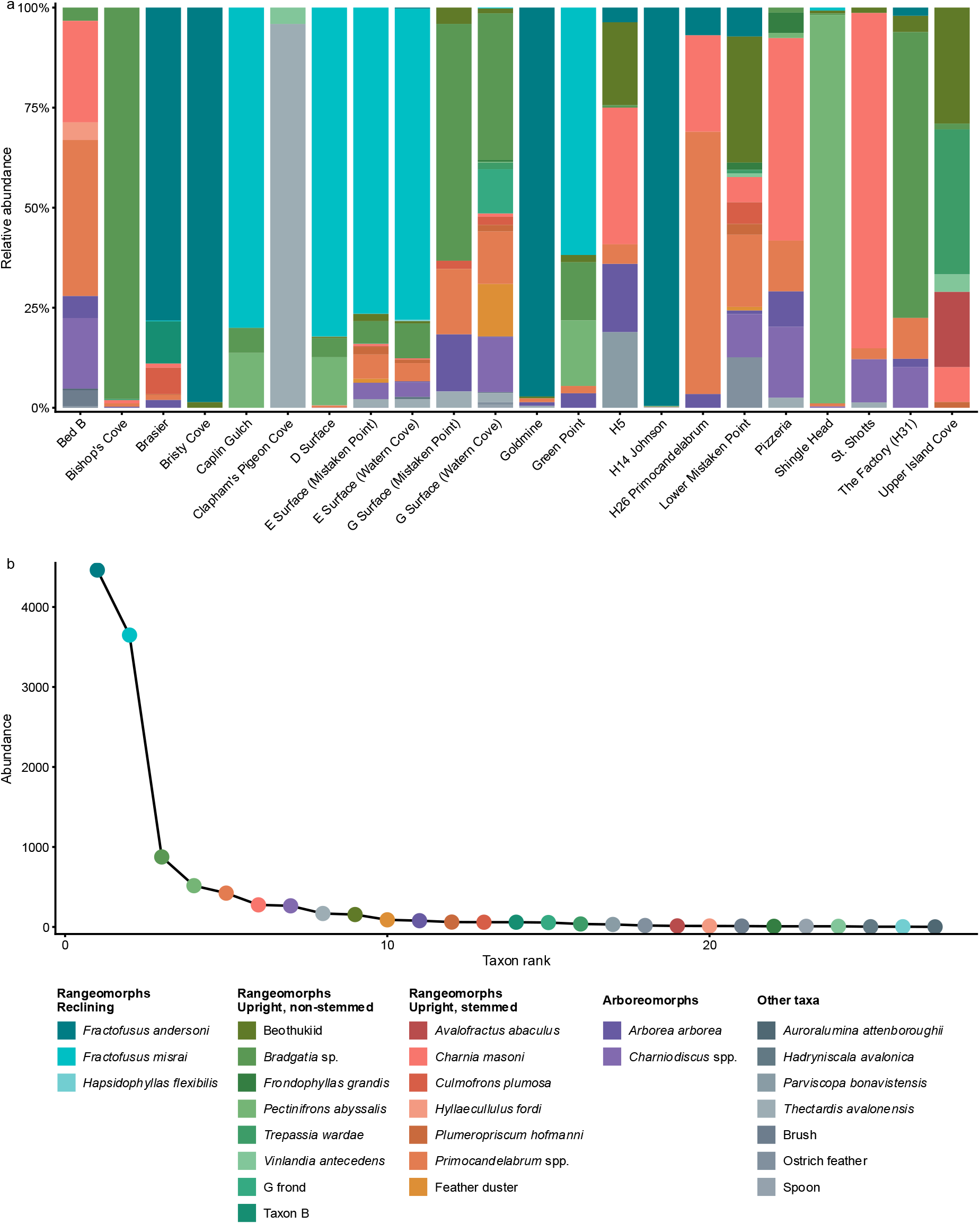
Taxonomic composition of Avalonian communities. (a) Relative abundance of taxa on each surface. (b) Pooled abundance–rank distribution across all surfaces.

### Community types and tiering

Hierarchical clustering identified compositional groups broadly consistent with the NMDS ordination, and mean silhouette width was highest at *k* = 7 (Fig. S2). Two closely associated clusters within this solution represented frond-dominated communities, one dominated largely by *Charnia* and *Primocandelabrum* and the other more compositionally mixed but with relatively high abundances of beothukiids (Fig. S2c). These two clusters were therefore combined into a single frond-dominated type, yielding four principal community types dominated respectively by *F. andersoni*; *F. misrai* and *Pectinifrons*; *Bradgatia*; and fronds, consistent with the community types described previously^17^. Clapham’s Pigeon Cove and Shingle Head were compositionally isolated in both analyses and were therefore retained as singleton categories; they occurred in the oldest and youngest formations represented, respectively (Fig. S2a,d; Table S1). The structure of the classification was retained after excluding E Surface at Watern Cove West and G Surface at Mistaken Point, the smaller E and G outcrops, respectively (Fig. S3), and under the finer-resolution taxonomic treatment (Fig. S4). Community-level DVS varied among the four principal community types, being generally highest in *F. andersoni*-dominated communities and lowest in frond-dominated communities, with intermediate values in the *F. misrai*–*Pectinifrons* and *Bradgatia* types (Kruskal–Wallis χ² = 8.55, *n* = 20, *p* = 0.036; Fig. S5).

### Sampling completeness

Spatially sampled taxon–area accumulation curves varied in the extent to which they approached an asymptote across the retrodeformed area of each surface (Fig. 2). Under the criterion that observed richness reached at least 80% of the fitted asymptote, 12 of the 22 surfaces were classified as saturated (Table S3). Saturated surfaces had higher median specimen numbers and retrodeformed areas than non-saturated surfaces, although neither difference was statistically significant (specimen number: Wilcoxon rank-sum *W* = 71.5, *n* = 22, *p* = 0.468; retrodeformed area: *W* = 76, *n* = 22, *p* = 0.307; Fig. S6). The finer-resolution taxonomic treatment produced broadly the same pattern, with only Pizzeria shifting to non-saturated as its observed richness fell just below 80% of the fitted asymptote (79.3%; Fig. S7; Table S4).

**Figure 2.**
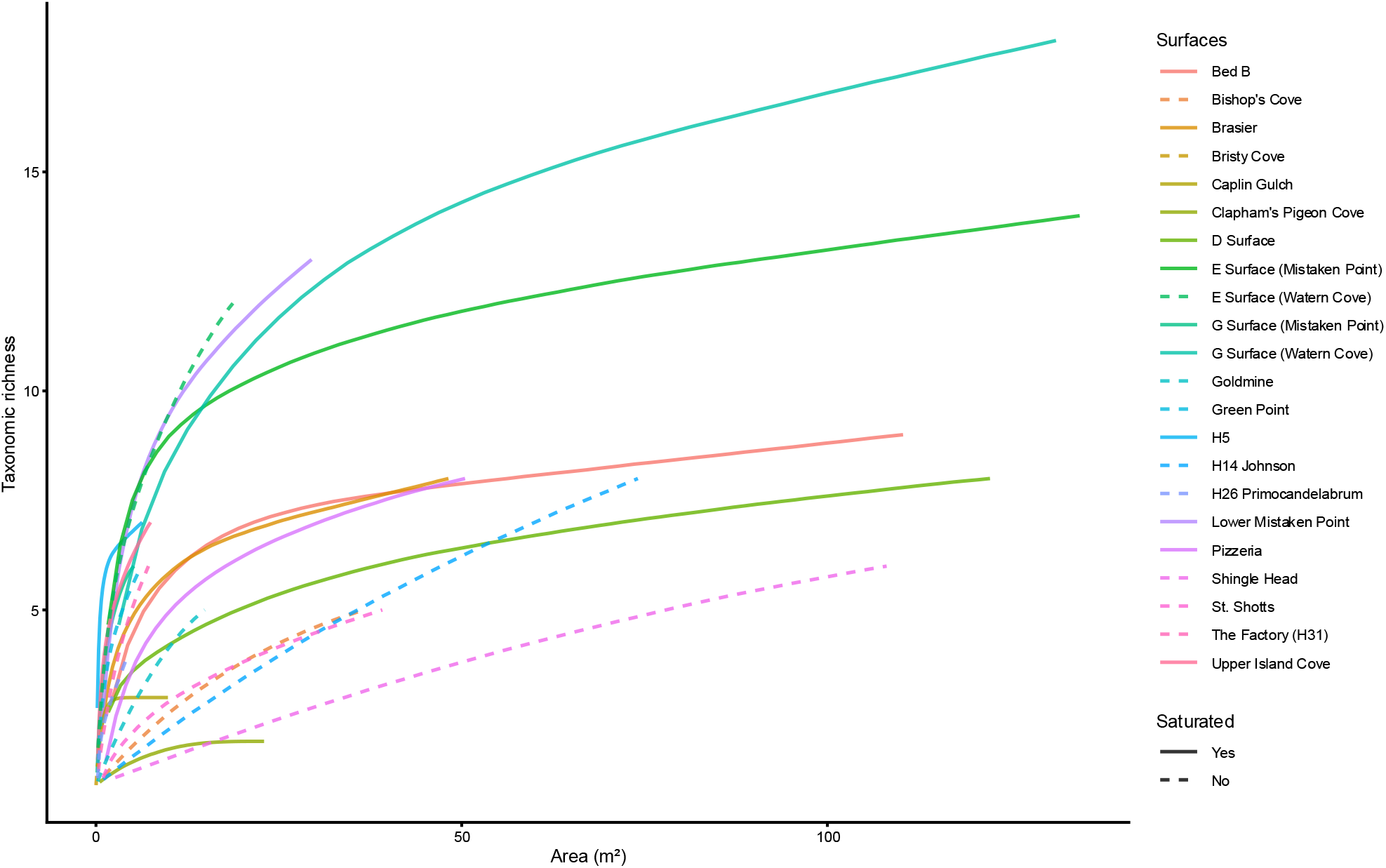
Taxon–area accumulation curves and sampling saturation across Avalonian fossiliferous surfaces. Model identities, fitted asymptotes and classifications are reported in Table S3; uncertainty around the accumulation curves is shown in Fig. S8.

### Diversity metrics

Among the four diversity metrics, observed and estimated Shannon and Simpson diversity were closely aligned across surfaces, with comparatively narrow uncertainty intervals (Fig. 3; Table S5). In contrast, extrapolated richness had wide upper intervals on several incompletely sampled surfaces, whereas evenness estimates propagated uncertainty from both richness and Shannon diversity. Given the greater uncertainty in asymptotic richness and evenness, subsequent candidate-correlate analyses focused on Shannon diversity, with the same analyses repeated using Simpson diversity. The finer-resolution treatment yielded higher observed and estimated richness on several surfaces (Fig. S9). Between the two taxonomic treatments, concordance was almost perfect for Simpson diversity (CCC = 0.998) and evenness (CCC = 0.999), substantial for Shannon diversity (CCC = 0.987 and 0.984), and poor to moderate for richness (CCC = 0.940 and 0.881; Fig. S10).

**Figure 3.**
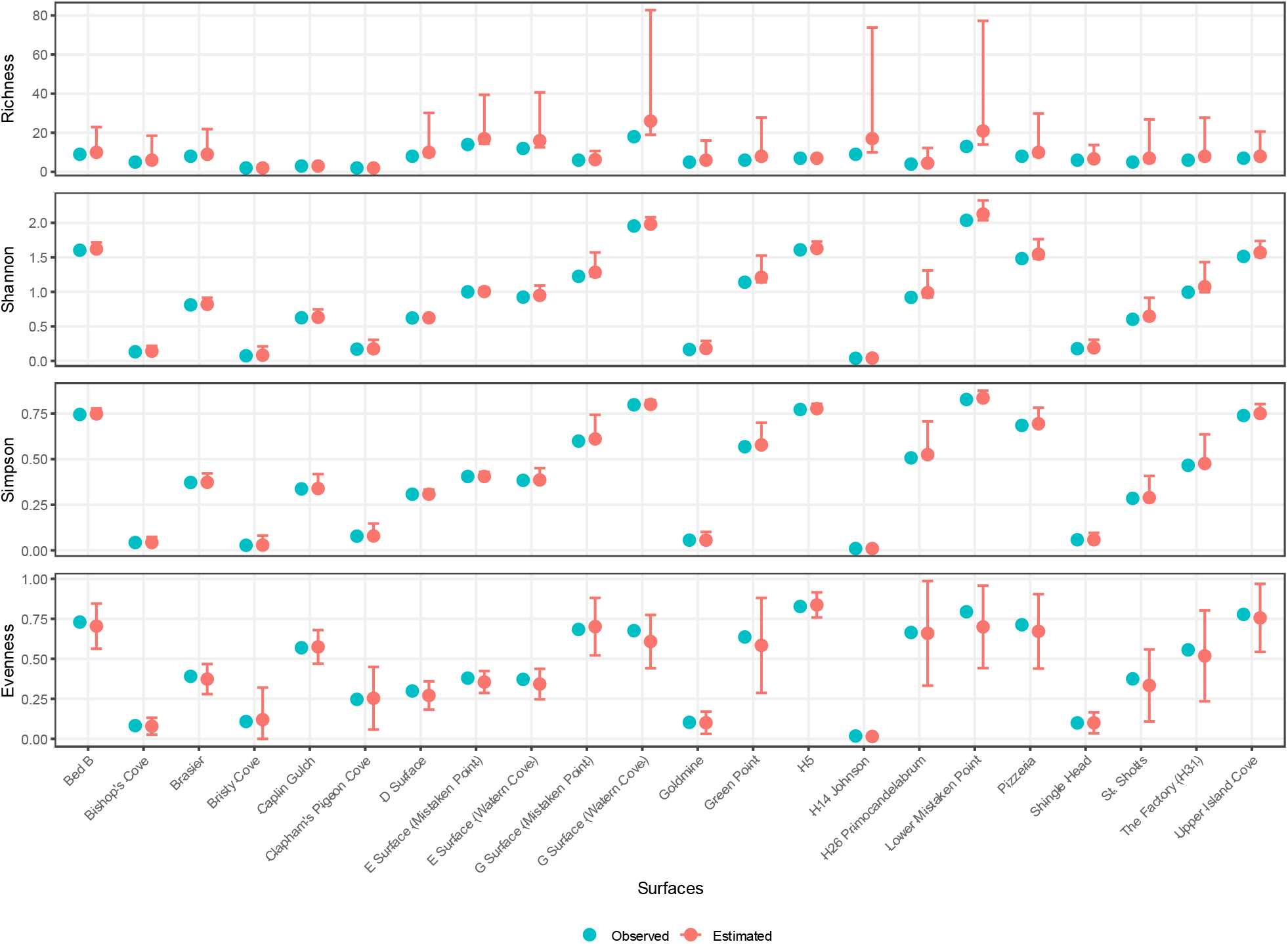
Observed and estimated diversity across Avalonian fossiliferous surfaces. The four rows show taxonomic richness, Shannon, Simpson and evenness, respectively. Vertical bars indicate 95% confidence intervals for estimated values. Numerical estimates and confidence intervals are reported in Table S5.

### Sampling effects

Shannon diversity showed no significant relationship with either specimen number or retrodeformed area when analysed as observed diversity among saturated surfaces, observed diversity across all surfaces, or estimated diversity across all surfaces (Fig. 4). The corresponding diversity–sampling regressions remained non-significant when the analyses were repeated using Simpson diversity, after excluding E Surface (Watern Cove West) and G Surface (Mistaken Point) outcrops to reduce potential non-independence from repeated sampling of the same stratigraphic horizons, or under the finer-resolution taxonomic treatment (Figs. S11–S13).

**Figure 4.**
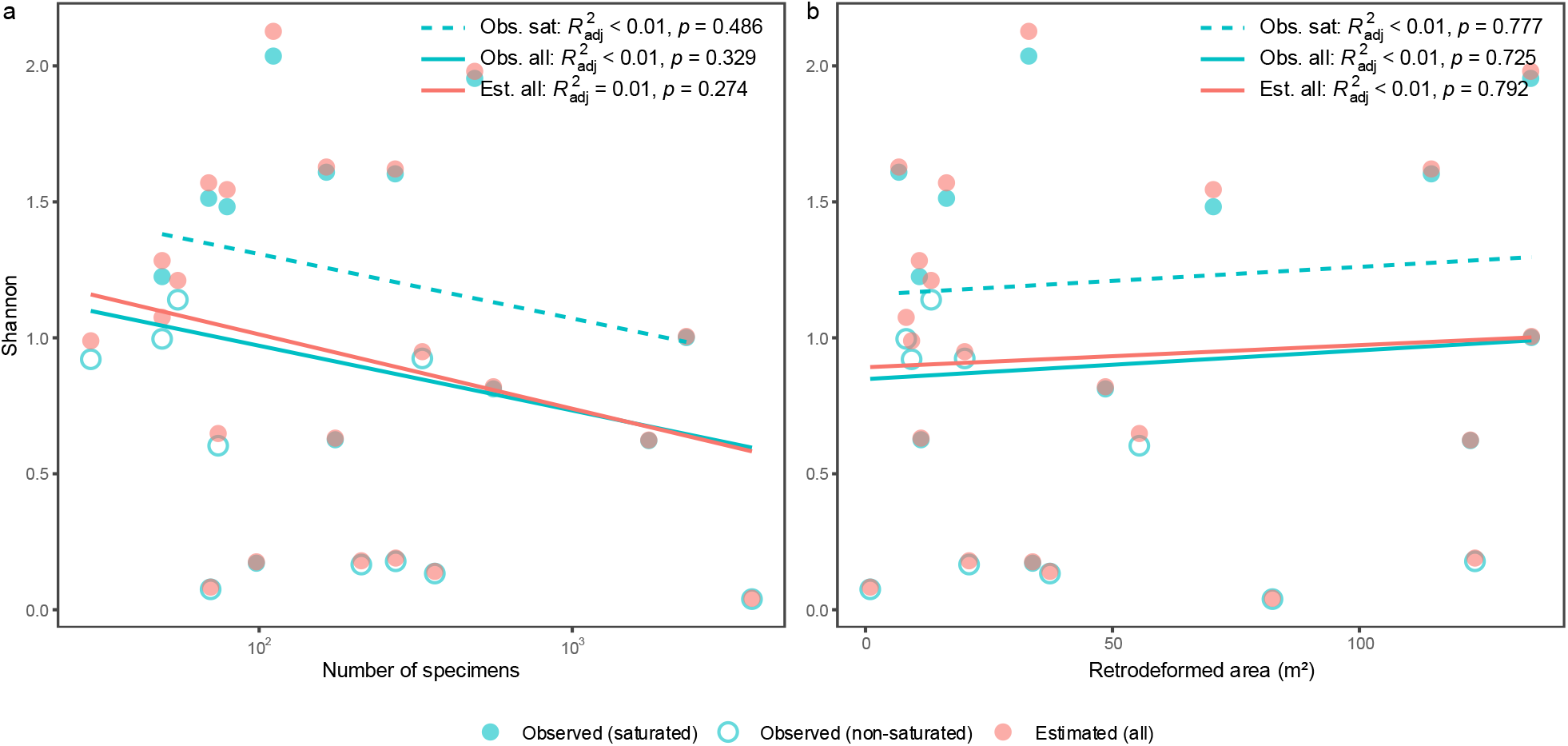
Relationships of Shannon diversity with sampling effort. Shannon diversity was analysed in relation to (a) number of specimens and (b) retrodeformed area, using observed values for saturated surfaces, observed values across all surfaces and estimated values across all surfaces. Adjusted *R*² and *p*-values for each regression are reported in each panel.

### Candidate correlates of diversity

Shannon diversity showed no clear association with locality or stratigraphic grouping. Among saturated surfaces, insufficient replication precluded a formal comparison among localities. Across all surfaces, observed diversity did not differ among Mistaken Point Ecological Reserve, Discovery Geopark and Spaniard’s Bay (Kruskal–Wallis χ² = 0.51, *n* = 21, *p* = 0.776), and estimated diversity likewise showed no significant difference (χ² = 0.32, *n* = 21, *p* = 0.85; Fig. 5a). At the level of the original stratigraphic units, only the Mistaken Point and Fermeuse formations contained sufficient surfaces for formal comparison, and neither observed nor estimated diversity differed significantly between them (both Wilcoxon rank-sum tests yielded *W* = 20, *n* = 12, *p* = 0.267; Fig. S14). When stratigraphically correlated units were combined, diversity also did not differ among formations for observed values across all surfaces (χ² = 1.89, *n* = 18, *p* = 0.39), or estimated values across all surfaces (χ² = 1.73, *n* = 18, *p* = 0.42; Fig. 5b).

**Figure 5.**
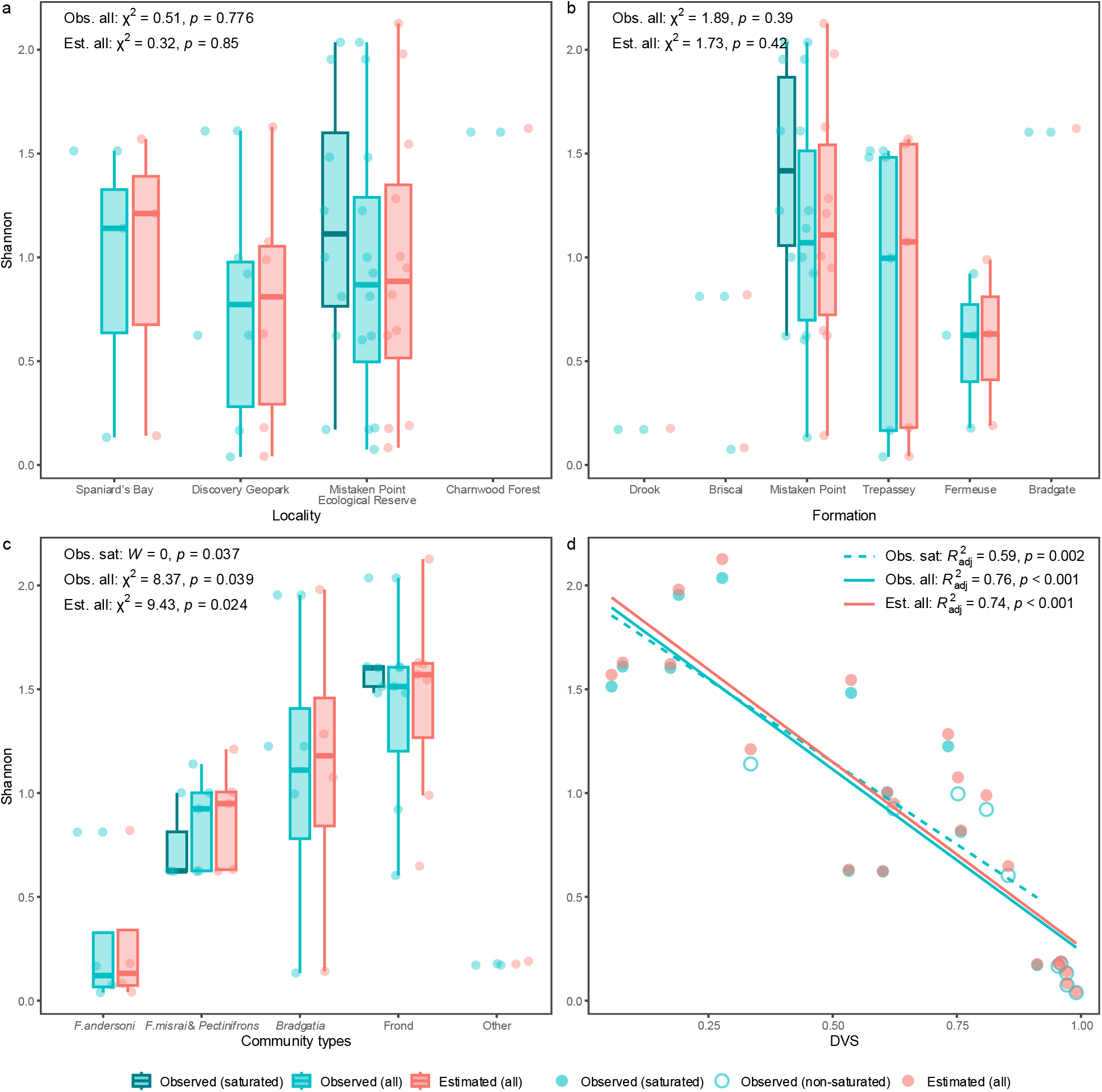
Relationships of Shannon diversity with candidate abiotic and biotic correlates. Shannon diversity was compared among (a) localities, (b) correlated formations and (c) community types, and analysed in relation to (d) community-level DVS, using observed values for saturated surfaces, observed values across all surfaces and estimated values across all surfaces. Categories represented by fewer than three surfaces were excluded from formal group comparisons. Wilcoxon rank-sum *W* and Kruskal–Wallis χ² are reported for categorical comparisons (a–c), and adjusted R² for linear regressions in (d), with corresponding *p*-values.

In contrast, both biotic correlates examined were associated with Shannon diversity. Among saturated surfaces, observed diversity was lower in *F. andersoni*-dominated than in upright frond-dominated communities (*W* = 0, *n* = 8, *p* = 0.037; Fig. 5c). Across all surfaces, both observed (χ² = 8.37, *n* = 20, *p* = 0.039) and estimated Shannon diversity (χ² = 9.43, *n* = 20, *p* = 0.024) differed among the four principal community types (Fig. 5c). Values increased broadly from *F. andersoni*-dominated communities, through the *F. misrai*–*Pectinifrons* and *Bradgatia*-dominated types, to upright frond-dominated communities (Fig. 5c). Shannon diversity was also strongly negatively associated with DVS across all three treatments (adj-*R*² = 0.59–0.76, all *p* ≤ 0.002; Fig. 5d).

Although DVS differed among community types (Fig. S5), the negative relationship between DVS and Shannon diversity was not solely attributable to between-type differences. In all three analyses, DVS-only models received stronger support than community-type-only models (Table S6). The negative trend was also broadly consistent within four principal community types (Fig. S15). The overall patterns were comparable for Simpson diversity and remained robust after excluding the two smaller outcrops of E and G surfaces and under the finer-resolution taxonomic treatment (Figs. S16–S18).

## Discussion

Avalonian fossiliferous surfaces exhibited pronounced heterogeneity in community composition, but most could nevertheless be organised into a small number of recurring groups. Of the 22 surfaces, 20 fell within four broad community types identified previously^17^, characterised by high relative abundances of *Fractofusus andersoni*, *F. misrai* and/or *Pectinifrons abyssalis*, *Bradgatia*, or a range of upright frondose taxa (e.g., *Charnia, Primocandelabrum*), and accompanied by differences in vertical organisation. The upright-frond type showed the greatest internal variation, with the clustering indicating several potentially overlapping subsets. The two remaining surfaces formed distinct configurations. Clapham’s Pigeon Cove comprised only *Thectardis avalonensis* and *Vinlandia avalonensis*, an unusual combination of taxa that were rare on most other surfaces, whereas Shingle Head was composed almost entirely of *Pectinifrons abyssalis* but lacked the substantial *F. misrai* component present in the other members of the broader *F. misrai*–*Pectinifrons* grouping. Their occurrence in the oldest and youngest formations sampled, respectively, raises the possibility that the margins of the sampled stratigraphic succession record phases of taxonomic turnover and community restructuring.

Each mapped fossiliferous surface represents an irregularly exposed window onto an underlying community, such that effective sampling varies among surfaces in mapped area, specimen number and the recovery of low-abundance taxa. Across the 22 surfaces, the sampled areas captured most of the taxonomic richness on 12 surfaces, whereas the remaining ten would likely have yielded additional taxa with further outcrop/sampling. Although the more completely sampled surfaces tended to contain more specimens and cover larger areas, neither difference was significant, indicating that taxonomic completeness was not a simple function of sampling effort alone.

Taxonomic completeness may therefore depend not only on the extent of fossiliferous bedding-plane exposure, but also on community structure: compositionally simpler assemblages, or those in which taxa are more evenly distributed across the mapped area (e.g. Caplin Gulch), may approach saturation within smaller areas, whereas spatially heterogeneous assemblages with a long tail of rare taxa (e.g. Pizzeria) may require broader coverage before richness approaches saturation^56^. Importantly, this assessment concerns the completeness of accurate taxon numbers and does not directly indicate how fully other ecological properties, including relative-abundance structure, spatial patterns or vertical organisation, were represented.

This uneven taxon recovery affected diversity metrics differently. When observed taxon-abundance data were extrapolated to estimate asymptotic diversity, richness estimates, and consequently the evenness estimates derived from them, remained relatively uncertain on incompletely sampled surfaces, whereas Shannon and Simpson estimates were comparatively stable. Variation in taxonomic resolution had a stronger effect on the number of recognised taxa (richness) than on the diversity indices. Across alternative treatments of taxa for which species-level resolution was uncertain or inconsistently available, Shannon, Simpson and evenness remained highly concordant across surfaces, whereas richness was understandably more sensitive to taxonomic resolution. Together with the lack of clear relationships between either Shannon or Simpson diversity and either specimen number or mapped area, these results support the use of Shannon and Simpson as relatively stable measures for cross-surface diversity comparisons under uneven sampling and taxonomic uncertainty^57,58^. At the same time, continued taxonomic revision and more consistent species-level identification, where possible, would refine richness estimates and enable more precise comparisons of relative abundance and community composition.

The absence of detected associations with locality or formation provides no evidence for strong broad-scale effects of environment, geography or time, but does not rule out subtler influences operating below the resolution of these categories. Locality provides only a coarse representation of separation among communities and may not map directly onto dispersal connectivity. Dispersal or metacommunity mixing may therefore have weakened locality-level differences, but this cannot be resolved using the present coarse categories^2,19^. Likewise, formation combines stratigraphic age with depositional and preservational context^15,35^, and correlations among regional stratigraphic units introduce additional uncertainty^31–33^. Geographic and stratigraphic replication was also uneven: although Bed B extended the dataset to Charnwood Forest, it was the only surface representing both that locality and the Bradgate Formation, limiting the detection of broader-scale contrasts. Abiotic influences may therefore have been weak, scale-dependent, or obscured by limited turnover across the sampled interval, short-lived ecological variation within formations, or environmental differences that did not align with formation boundaries^17^. Distinguishing among these possibilities will require higher-resolution palaeoenvironmental information, including more direct sedimentological or geochemical covariates at the bed scale, together with greater replication across underrepresented geographic and stratigraphic settings where suitable surfaces are available.

Diversity varied systematically with recurring patterns of community composition, increasing from lower-diversity *Fractofusus*-rich communities through *Bradgatia*-dominated communities to higher-diversity upright-frond communities. These community types were organised around different dominant taxa and spanned multiple successional stages rather than representing successive phases of a single developmental sequence^17^. Therefore, diversity differences among types may reflect differences in community structure, whereas variation within types may partly reflect differences in community maturity. Vertical structure showed a stronger and more continuous association with diversity: diversity declined as DVS increased, indicating that the most diverse surfaces were characterised by greater overlap in vertical resource-acquisition space rather than stronger tiering. This pattern is more consistent with equalising processes that reduced dominance or fitness inequalities than with stabilising coexistence through stronger vertical niche partitioning^23,59,60^.

Community composition and vertical structure are closely linked aspects of community organisation, making their associations with diversity difficult to separate completely^17^. Distinguishing the processes underlying these patterns will therefore require considering how multiple aspects of biotic organisation interact rather than attributing diversity variation to a single mechanism. Nevertheless, the negative relationship between diversity and DVS was retained within each of the four principal community types. Model comparisons further indicated that community type and DVS captured related but not entirely interchangeable components of diversity variation. Their combined pattern was particularly apparent in the contrast between *Fractofusus*-rich and upright-frond communities: the former generally occupied a lower and narrower range of diversity, whereas the latter encompassed higher and more variable diversity. This contrast may indicate a more constrained set of realised community configurations in *Fractofusus*-rich assemblages and greater structural flexibility among upright-frond assemblages. However, high DVS on low-diversity surfaces may also partly reflect strong dominance by reclining taxa and the pronounced contrast between reclining and upright body plans, potentially because reproductive strategies such as prolific stoloniferous growth can alter competition dynamics rather than stabilising niche partitioning alone^17,24^. Resolving the relative contributions and interactions of community composition and vertical structure, as well as incorporating additional biotic and abiotic factors, will require greater replication across surfaces sufficient to support multivariable models containing both additive and interaction terms.

## Conclusions

Together, these analyses demonstrate both the value and the limitations of Avalon fossiliferous surfaces for comparative palaeoecology. Although taxon sampling was incomplete on some surfaces, Shannon and Simpson diversity were comparatively robust to uneven sampling and differences of taxonomic resolution, supporting cross-surface comparisons of abundance-weighted alpha diversity. These comparisons consistently linked diversity to recurring patterns of community composition and niche partitioning, whereas no association with age and environment (formation) or biogeography (locality) was detected. Avalon diversity is therefore best interpreted as covarying with both community composition and emergent vertical organisation, rather than with abiotic setting alone or any single ecological driver.

## Ethics

The Parks and Natural Areas Division, Department of Environment and Conservation, Government of Newfoundland and Labrador, provided permits to conduct research within the Mistaken Point Ecological Reserve in 2010, 2016–2019 and 2021–2023. Access to Mistaken Point Ecological Reserve is by scientific research permit only. Fossil surfaces on the Bonavista Peninsula and in the Spaniard’s Bay area are protected under Regulation 67/11 of the Historic Resources Act 2011, and fieldwork in the Discovery Geopark and Spaniard’s Bay areas was conducted under permits issued by the Government of Newfoundland and Labrador. Bed B forms part of a designated Site of Special Scientific Interest protected by law and administered by Natural England. No fossil specimens were removed as part of the present study.

## Data availability

The data required to reproduce the results of this study will be made public upon publication. Due to geoconservation concerns, data from Bed B cannot be made publicly available and will be subject to restricted access.

## Code availability

The code required to reproduce the results of this study will be made public upon publication.

## Acknowledgements

We thank Euan Furness for helpful discussions; Mark King, Richard Thomas and Edith Samson for their logistical support. N.P.S. was supported by a School of Biological Sciences Balfour Fund, University of Cambridge, and by the Roger Evans and Aey Phanachet Research Fellow in Symbiosis and Co-Evolution, King’s College, University of Cambridge. E.G.M. was supported by a Natural Environment Research Council Independent Research Fellowship NE/S014756/1.

## Author contributions

The study was conceptualised by all authors. Data processing and statistical analyses were conducted by S.C., N.P.S. and E.G.M. The project was supervised by A.M. and E.G.M. The original draft was written by S.C. and N.P.S. All authors contributed to revising and editing the final manuscript.

## Supplementary Information

**Figure S1.**
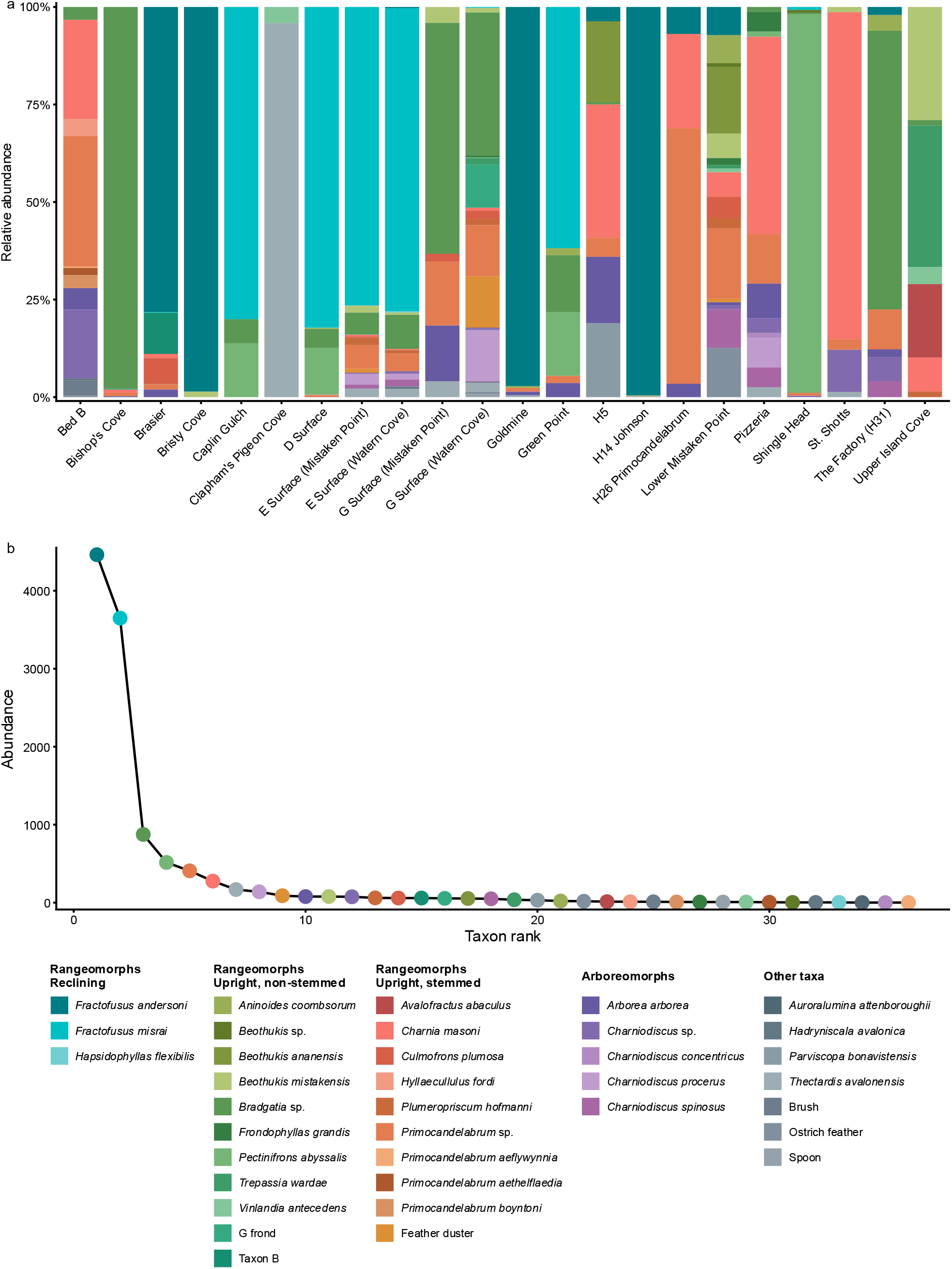
Taxonomic composition of Avalonian communities under the finer-resolution taxonomic treatment. (a) Relative abundance of taxa on each surface. (b) Pooled abundance–rank distribution across all surfaces. Corresponds to Fig. 1, with specimens within the “beothukiid”, *Charniodiscus* and *Primocandelabrum* groupings resolved to species level where possible.

**Figure S2.**
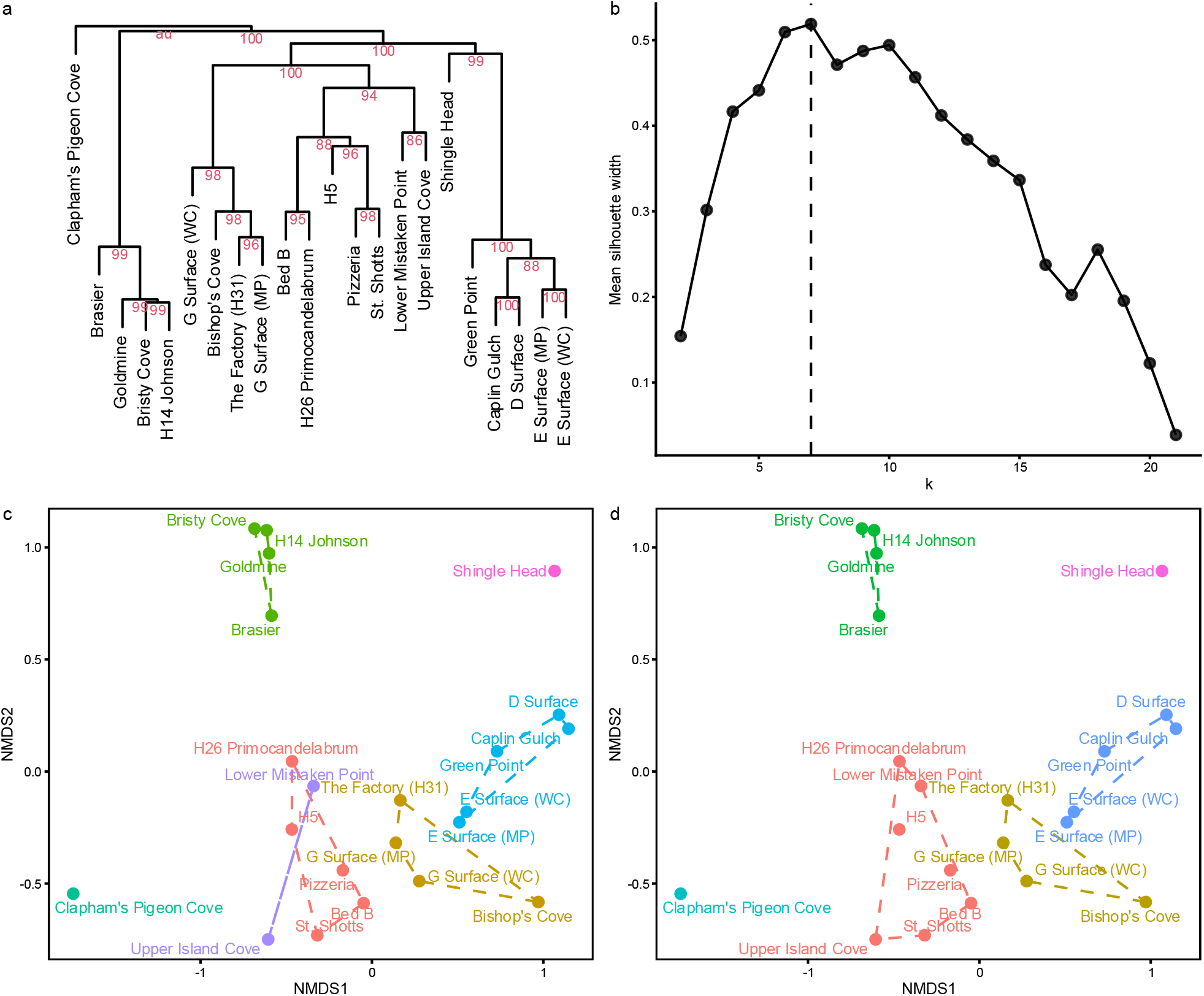
Community classification based on hierarchical clustering and NMDS ordination. (a) UPGMA clustering of Bray–Curtis dissimilarities among surfaces, with approximately unbiased bootstrap support values shown at internal nodes. (b) Mean silhouette width across candidate numbers of clusters, with the maximum at *k* = 7 indicated by the dashed line. (c) Two-dimensional NMDS ordination (stress = 0.136), grouped according to the silhouette-optimal clustering solution (*k* = 7). (d) Final community classification after combining the two closely overlapping frond-dominated clusters (*k* = 6), yielding four principal community types and two singleton categories. WC = Watern Cove; MP = Mistaken Point.

**Figure S3.**
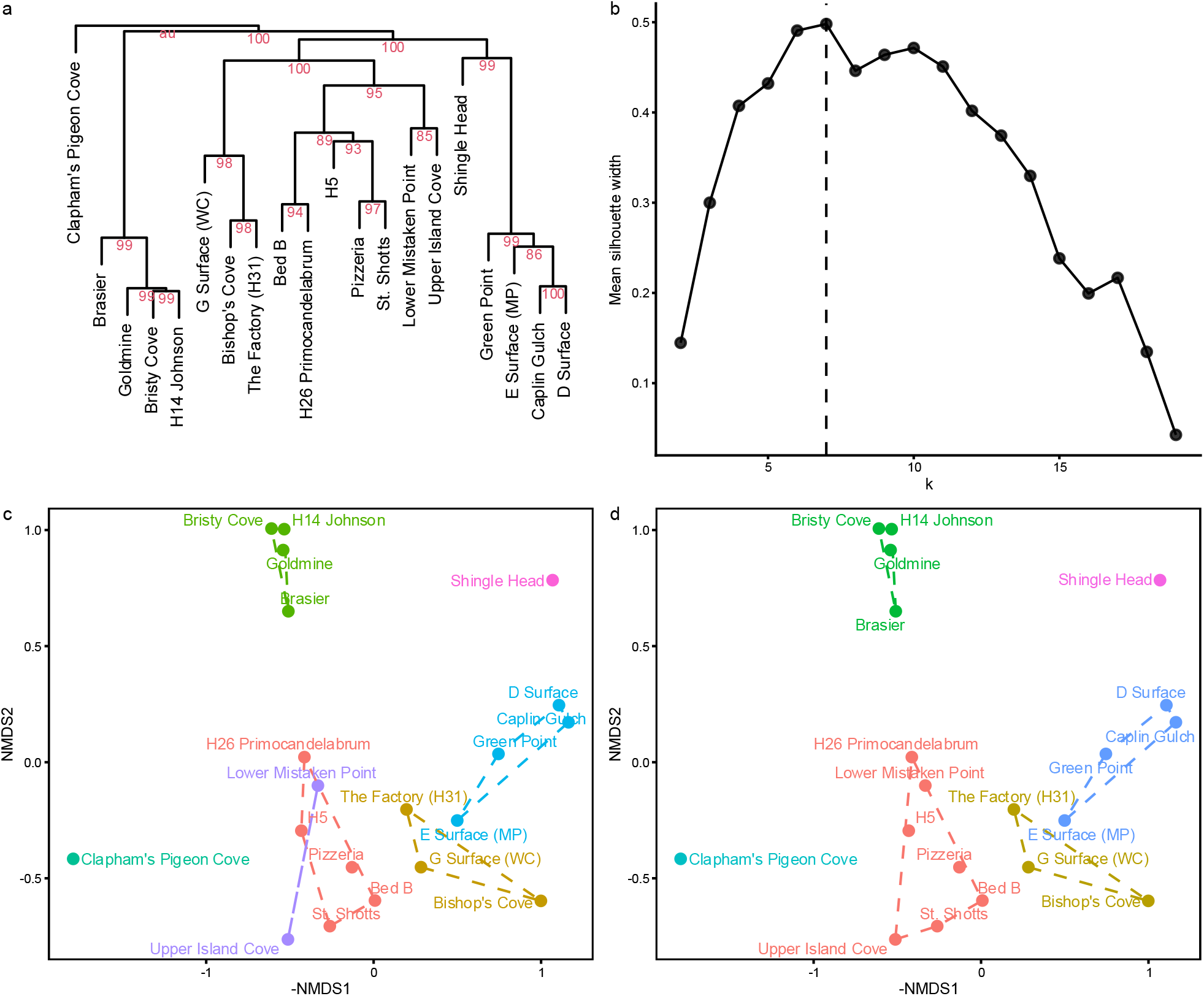
Community classification based on hierarchical clustering and NMDS ordination after excluding the smaller duplicated outcrops. (a) UPGMA clustering of Bray–Curtis dissimilarities among surfaces, with approximately unbiased bootstrap support values shown at internal nodes. (b) Mean silhouette width across candidate numbers of clusters, with the maximum at *k* = 7 indicated by the dashed line. (c) Two-dimensional NMDS ordination (stress = 0.124), grouped according to the silhouette-optimal clustering solution (*k* = 7). (d) Final community classification after combining the two closely overlapping frond-dominated clusters (*k* = 6), yielding four principal community types and two singleton categories. Corresponds to Fig. S2 after excluding E Surface (Watern Cove) and G Surface (Mistaken Point). WC = Watern Cove; MP = Mistaken Point.

**Figure S4.**
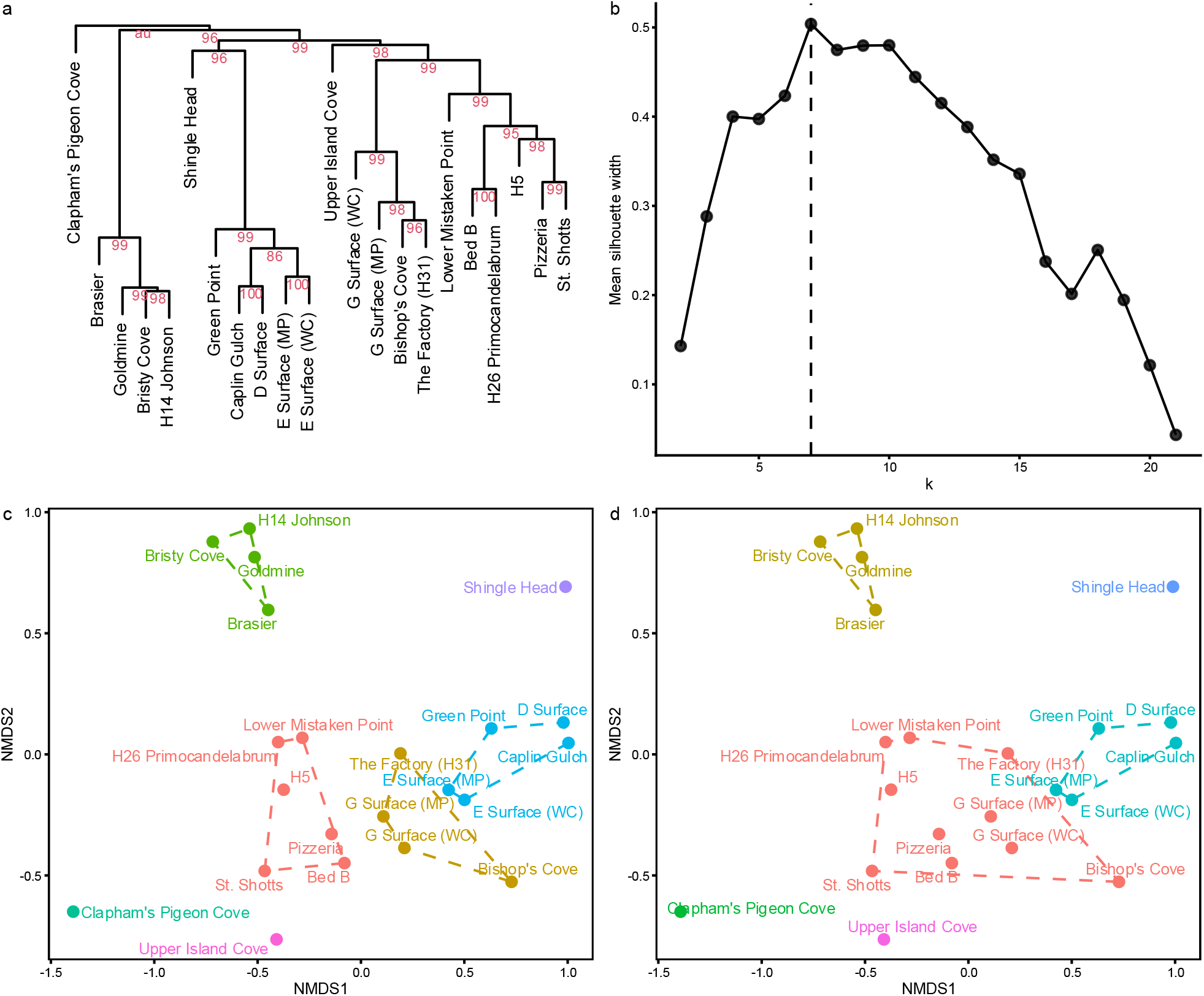
Community classification based on hierarchical clustering and NMDS ordination under the finer-resolution taxonomic treatment. (a) UPGMA clustering of Bray–Curtis dissimilarities among surfaces, with approximately unbiased bootstrap support values shown at internal nodes. (b) Mean silhouette width across candidate numbers of clusters, with the maximum at *k* = 7 indicated by the dashed line. (c) Two-dimensional NMDS ordination (stress = 0.139), grouped according to the silhouette-optimal clustering solution (*k* = 7), which recovered four principal groups and three singleton surfaces. (d) The *k* = 6 solution shown for comparison with the main taxonomic treatment; under this solution, the *Bradgatia*-dominated and frond-dominated groups were combined. Corresponds to Fig. S2, with specimens within the “beothukiid”, *Charniodiscus* and *Primocandelabrum* groupings resolved to species level where possible. WC = Watern Cove; MP = Mistaken Point.

**Figure S5.**
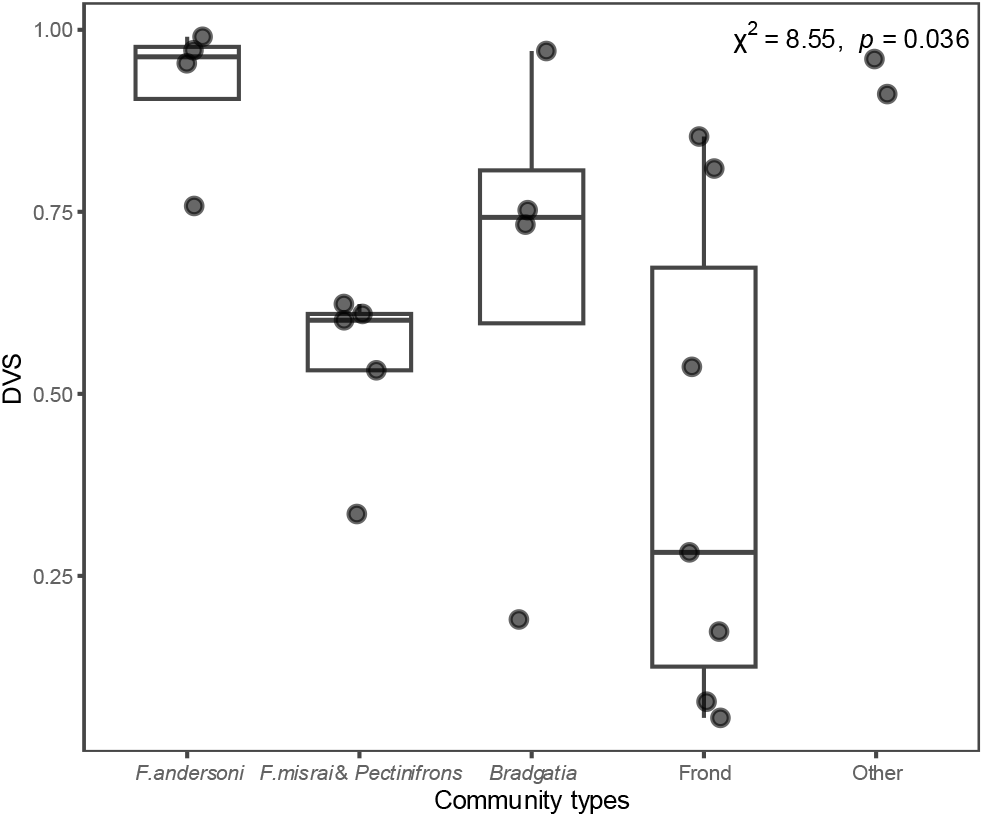
Community-level DVS among community types. Community-level DVS was compared among the four principal community types. The two singleton categories are shown together as Other for visual reference but were excluded from the Kruskal–Wallis test and are therefore displayed without a boxplot. Kruskal–Wallis χ² and *p*-value are reported in the panel.

**Figure S6.**
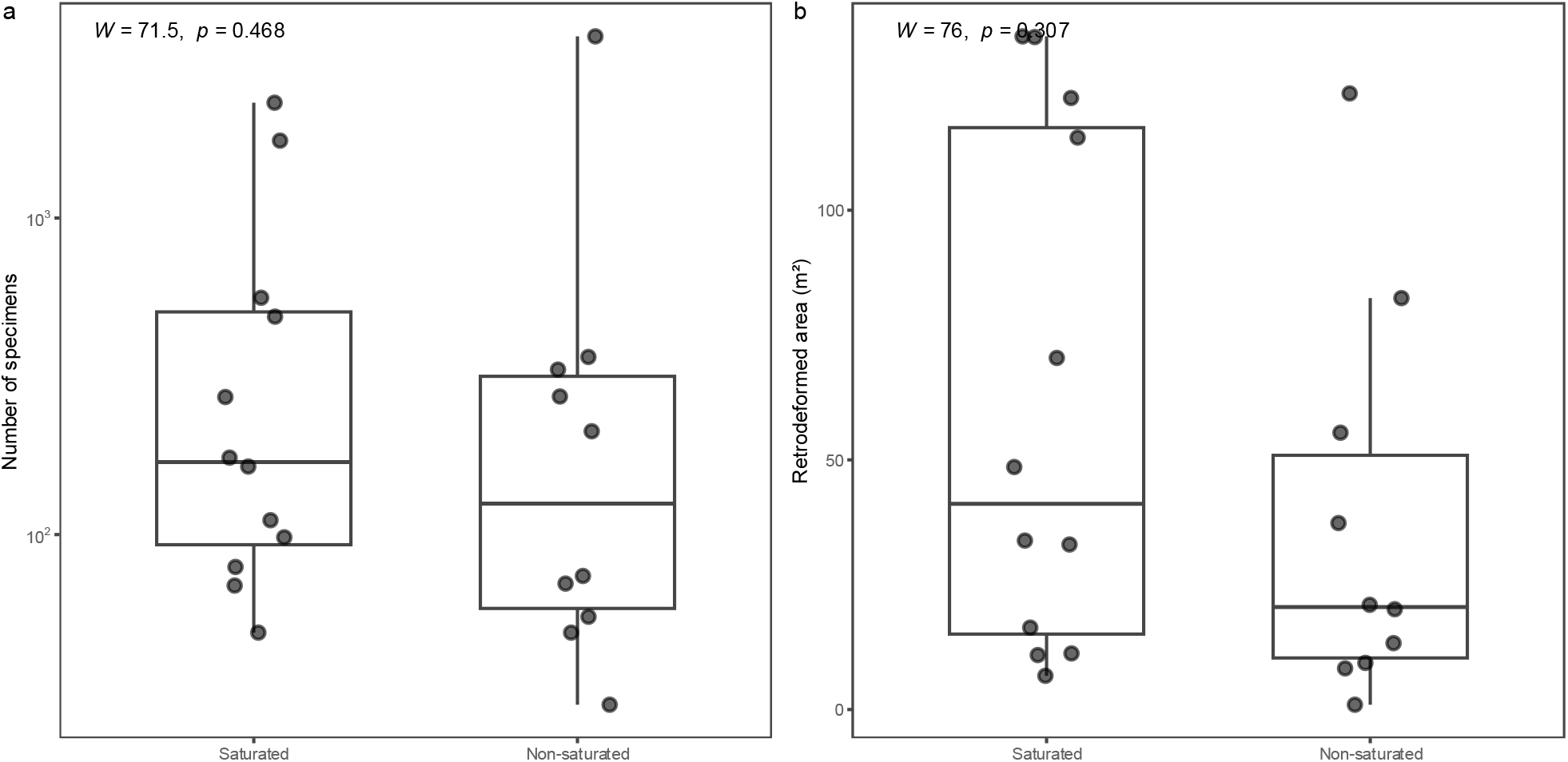
Sampling effort for saturated and non-saturated surfaces. The number of specimens (a) and retrodeformed area (b) were compared between saturated and non-saturated surfaces. Wilcoxon rank-sum *W* and *p*-values are reported in each panel.

**Figure S7.**
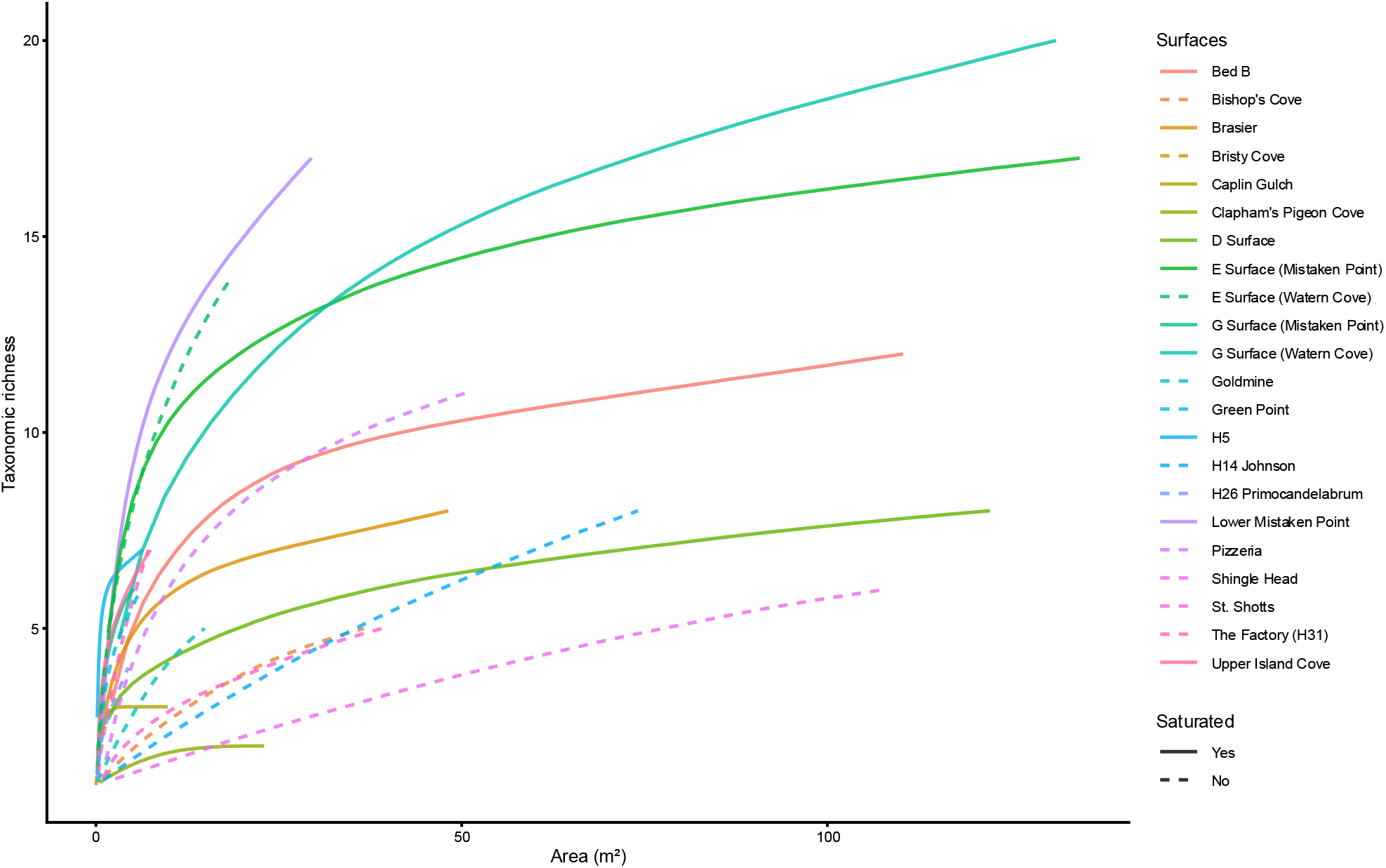
Taxon–area accumulation curves and sampling saturation across Avalonian fossiliferous surfaces under the finer-resolution taxonomic treatment. Corresponds to Fig. 2, with specimens within the “beothukiid”, *Charniodiscus* and *Primocandelabrum* groupings resolved to species level where possible. Model identities, fitted asymptotes and saturation classifications are reported in Table S4.

**Figure S8.**
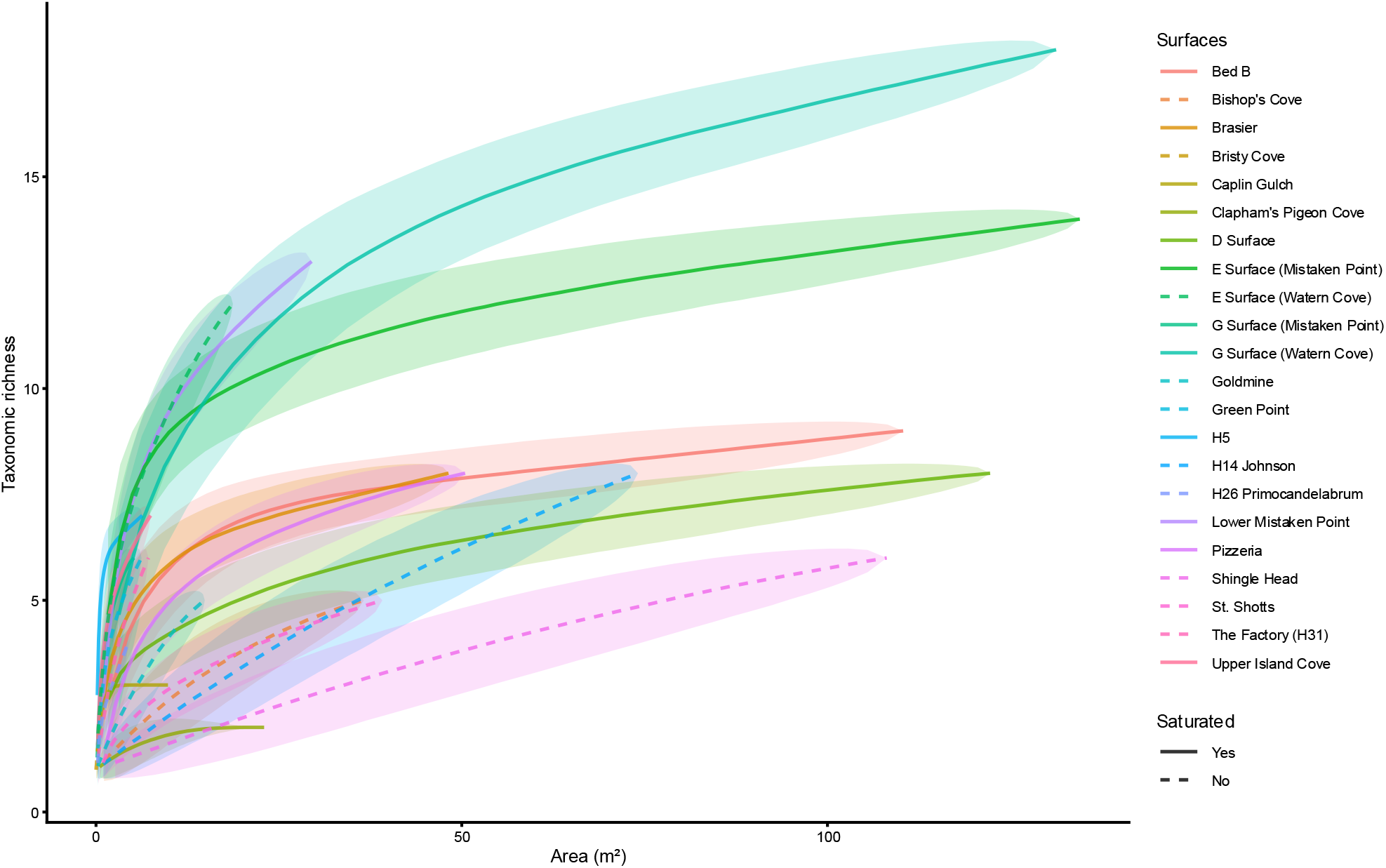
Taxon–area accumulation curves and sampling saturation across Avalonian fossiliferous surfaces. As in Fig. 2, with shaded ribbons indicating ±1 standard deviation across 10,000 permutations.

**Figure S9.**
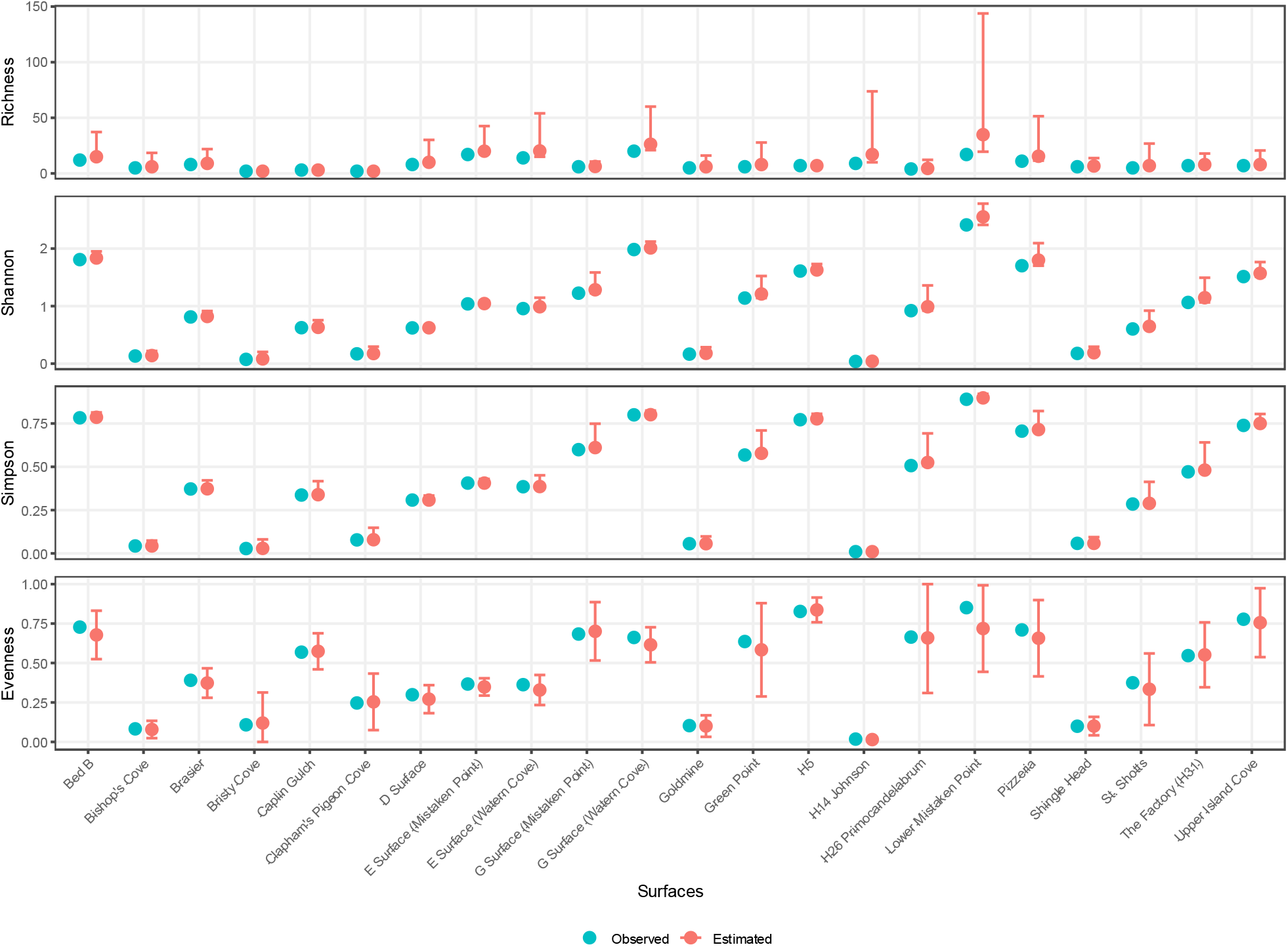
Observed and estimated diversity across Avalonian fossiliferous surfaces under the finer-resolution taxonomic treatment. The four rows show taxonomic richness, Shannon, Simpson and evenness, respectively. Vertical bars indicate 95% confidence intervals for estimated values. Corresponds to Fig. 3, with specimens within the “beothukiid”, *Charniodiscus* and *Primocandelabrum* groupings resolved to species level where possible.

**Figure S10.**
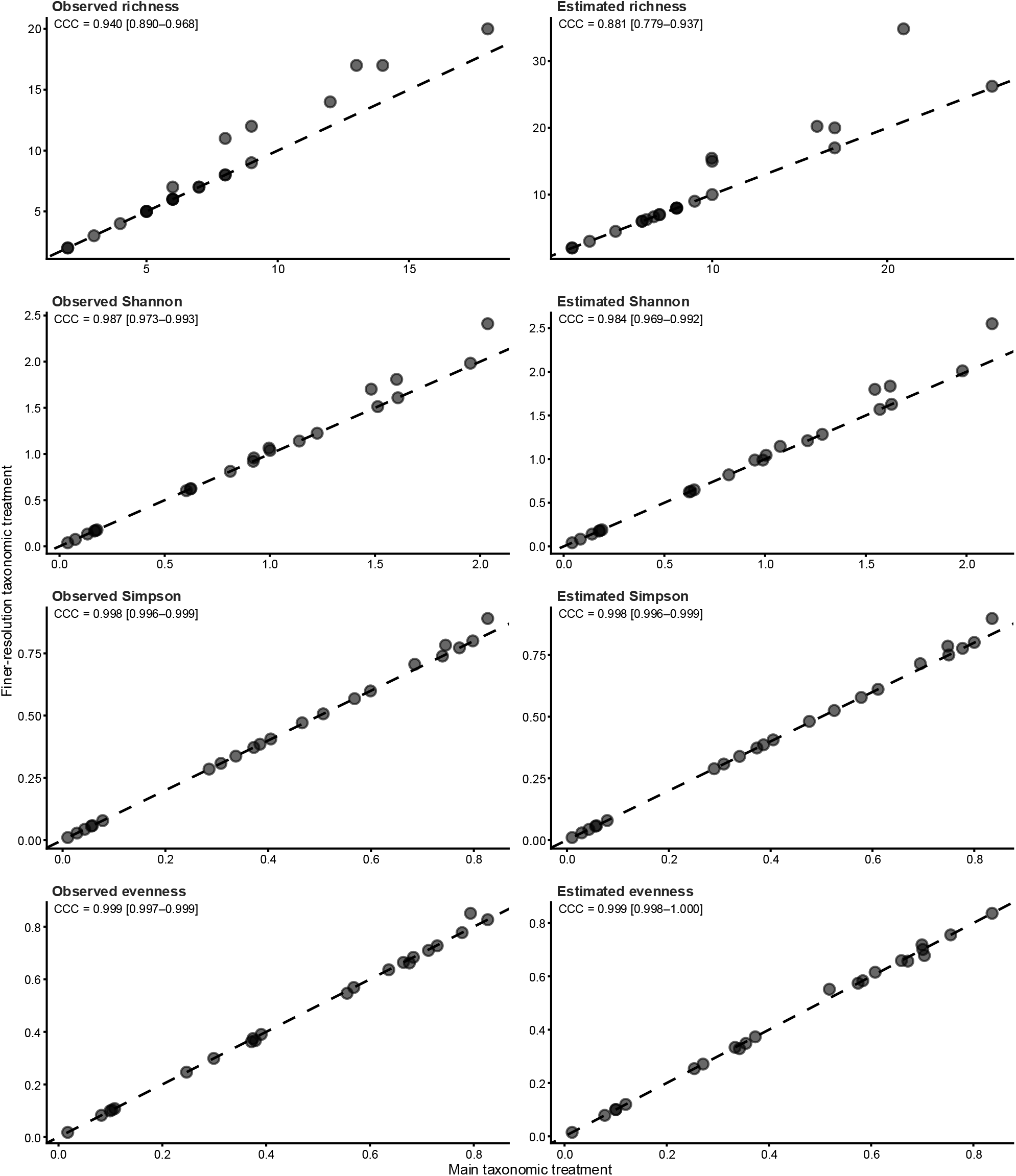
Agreement in diversity estimates between the main and finer-resolution taxonomic treatments. Panels compare observed and estimated taxonomic richness, Shannon, Simpson and evenness under the main taxonomic treatment (x-axis) and the finer-resolution sensitivity treatment (y-axis). Dashed lines indicate the 1:1 line of perfect concordance. Lin’s concordance correlation coefficient (CCC), with 95% confidence intervals, is reported in each panel.

**Figure S11.**
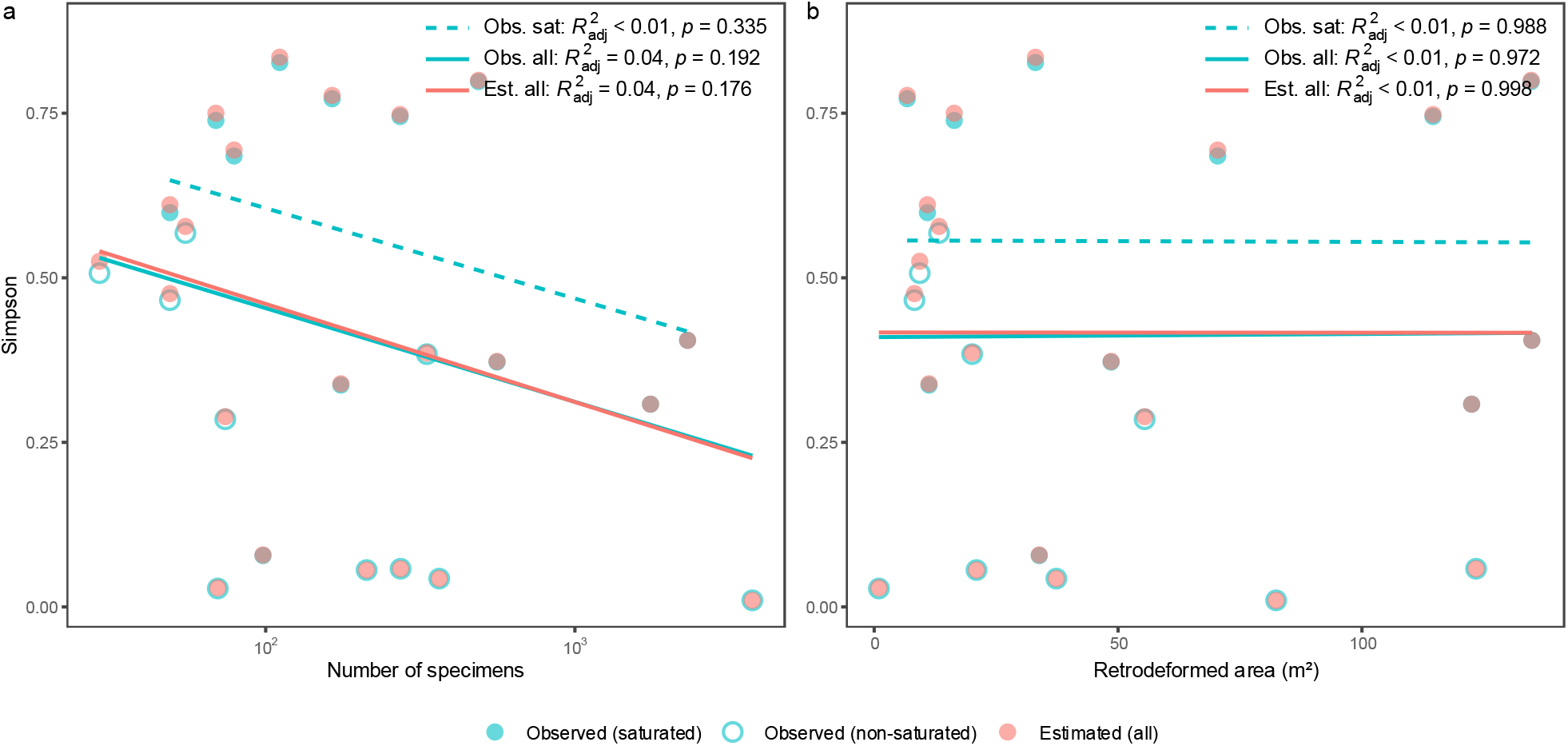
Relationships of Simpson diversity with sampling effort. Simpson diversity was analysed in relation to (a) number of specimens and (b) retrodeformed area, using observed values for saturated surfaces, observed values across all surfaces and estimated values across all surfaces. Adjusted *R*² and *p*-values for each regression are reported in each panel. Corresponds to Fig. 4, but using Simpson diversity.

**Figure S12.**
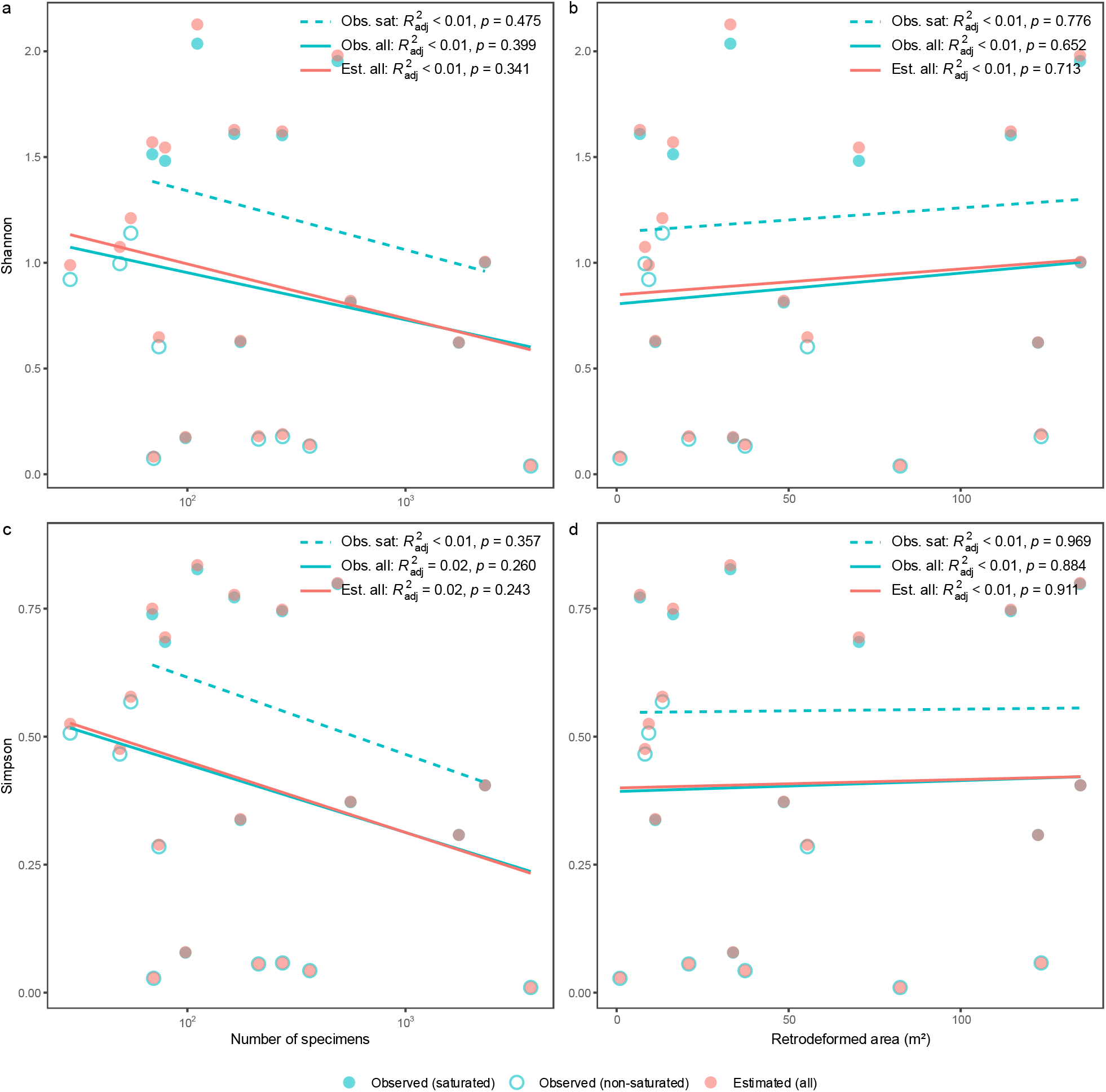
Relationships of Shannon and Simpson diversity with sampling effort after excluding the smaller duplicated outcrops. Shannon diversity (a,b) and Simpson diversity (c,d) were analysed in relation to number of specimens (a,c) and retrodeformed area (b,d), using observed values for saturated surfaces, observed values across all surfaces and estimated values across all surfaces. Adjusted *R*² and *p*-values for each regression are reported in each panel. Corresponds to Fig. 4 after excluding E Surface (Watern Cove) and G Surface (Mistaken Point).

**Figure S13.**
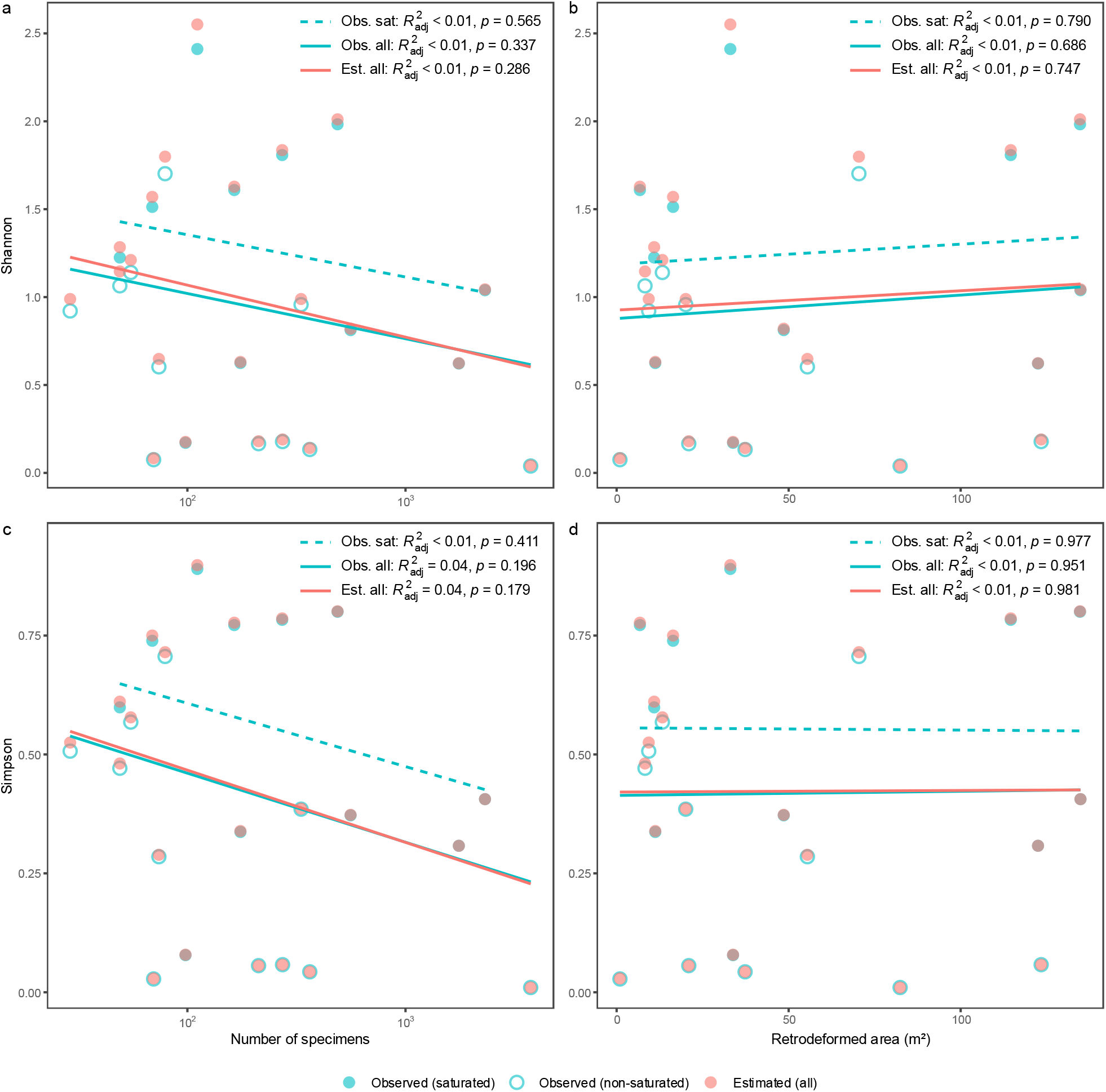
Relationships of Shannon and Simpson diversity with sampling effort under the finer-resolution taxonomic treatment. Shannon diversity (a,b) and Simpson diversity (c,d) were analysed in relation to number of specimens (a,c) and retrodeformed area (b,d), using observed values for saturated surfaces, observed values across all surfaces and estimated values across all surfaces. Adjusted *R*² and *p*-values for each regression are reported in each panel. Corresponds to Fig. 4, with specimens within the “beothukiid”, *Charniodiscus* and *Primocandelabrum* groupings resolved to species level where possible.

**Figure S14.**
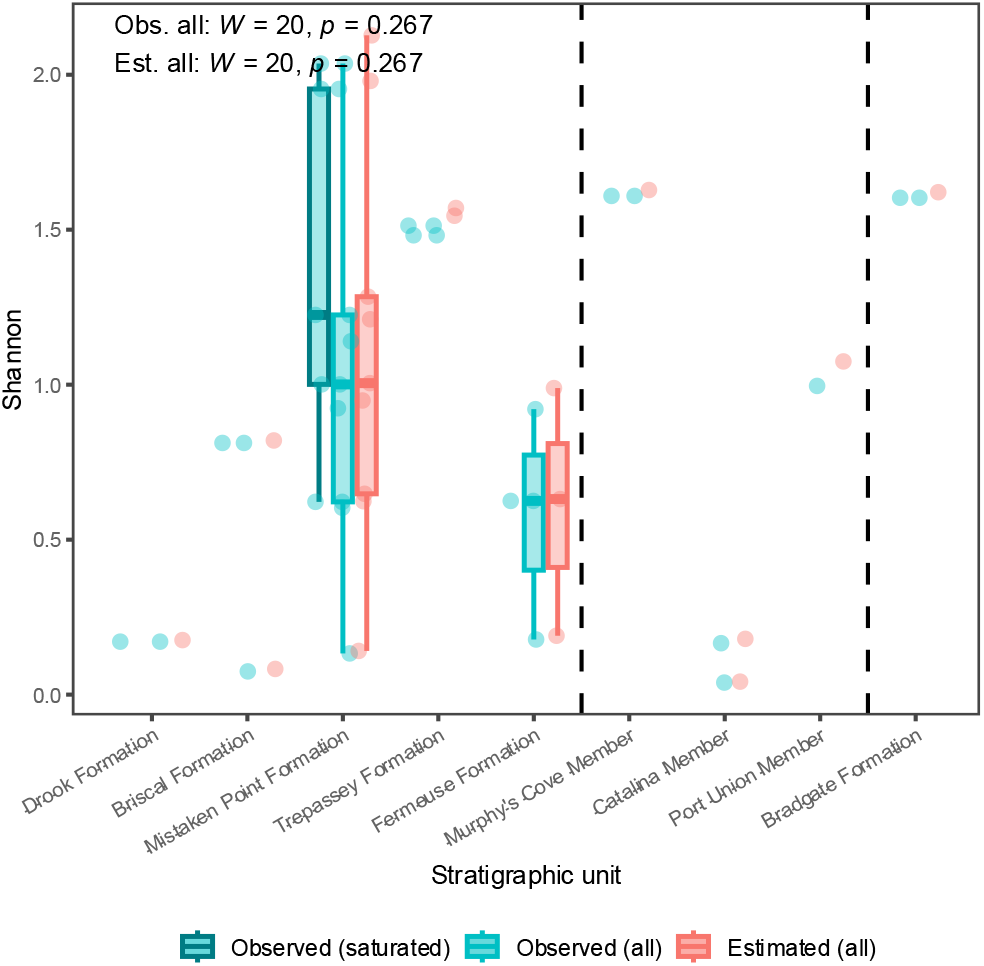
Relationships of Shannon diversity with the original stratigraphic units. Shannon diversity was compared among stratigraphic units using observed values for saturated surfaces, observed values across all surfaces and estimated values across all surfaces. Stratigraphic units are grouped regionally and ordered stratigraphically within each group; dashed vertical lines separate the combined Mistaken Point Ecological Reserve and Spaniard’s Bay units on the left, Discovery Geopark in the middle, and Charnwood Forest on the right. Categories represented by fewer than three surfaces were excluded from formal group comparisons. Wilcoxon rank-sum *W* and *p*-values are reported in the panel. Corresponds to Fig. 5b, but using the original, uncorrelated stratigraphic units.

**Figure S15.**
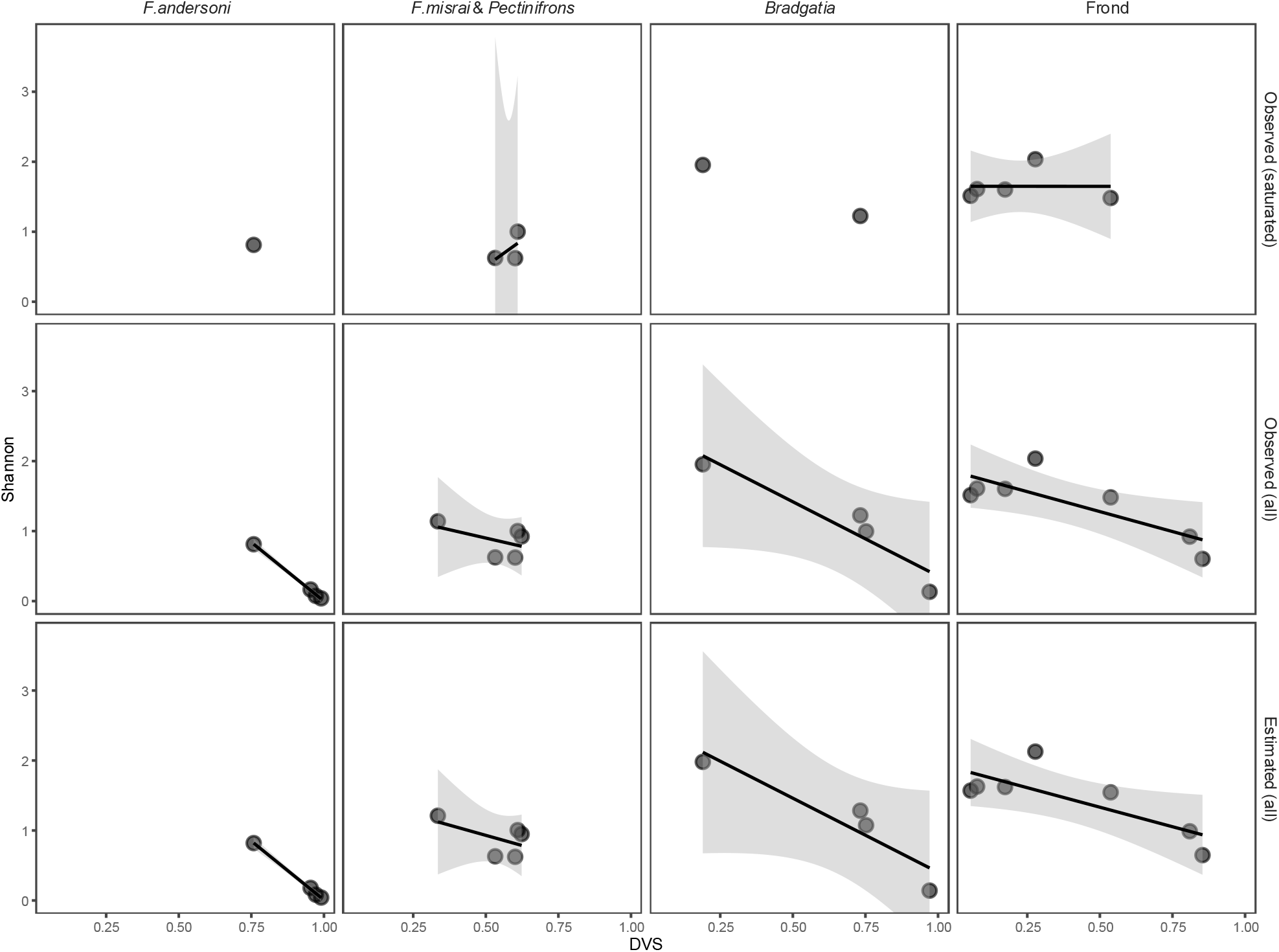
Relationships of Shannon diversity with DVS across the four principal community types. Relationships are shown for observed diversity among saturated surfaces, observed diversity across all surfaces and estimated diversity across all surfaces. Points represent individual surfaces, with fitted linear regressions and 95% confidence intervals shown where at least three surfaces were available.

**Figure S16.**
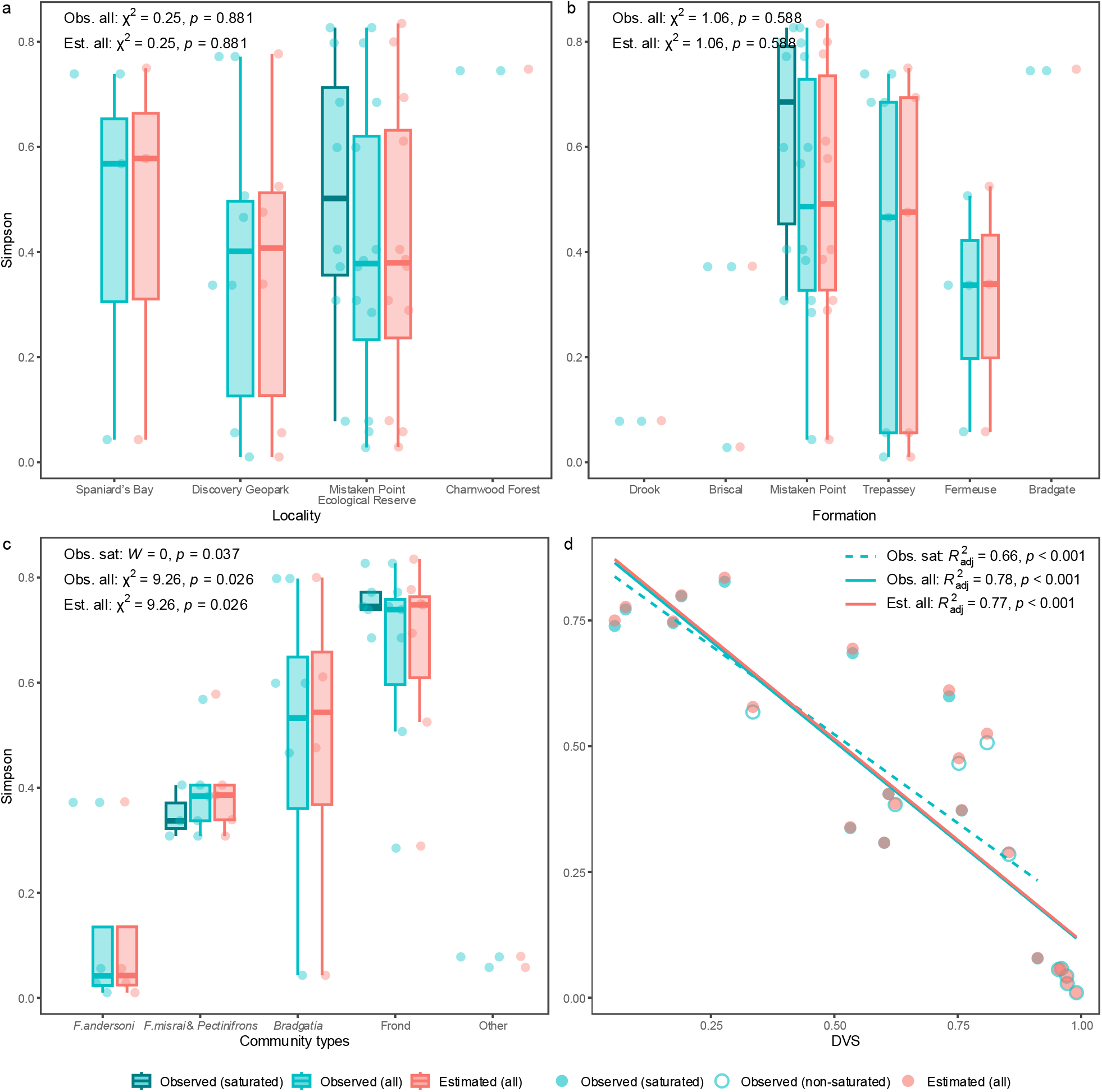
Relationships of Simpson diversity with candidate abiotic and biotic correlates. Simpson diversity was compared among (a) localities, (b) correlated formations and (c) community types, and analysed in relation to (d) community-level DVS, using observed values for saturated surfaces, observed values across all surfaces and estimated values across all surfaces. Categories represented by fewer than three surfaces were excluded from formal group comparisons. Wilcoxon rank-sum *W* and Kruskal–Wallis χ² are reported for categorical comparisons (a–c), and adjusted *R*² for linear regressions in (d), with corresponding *p*-values. Corresponds to Fig. 5, but using Simpson diversity.

**Figure S17.**
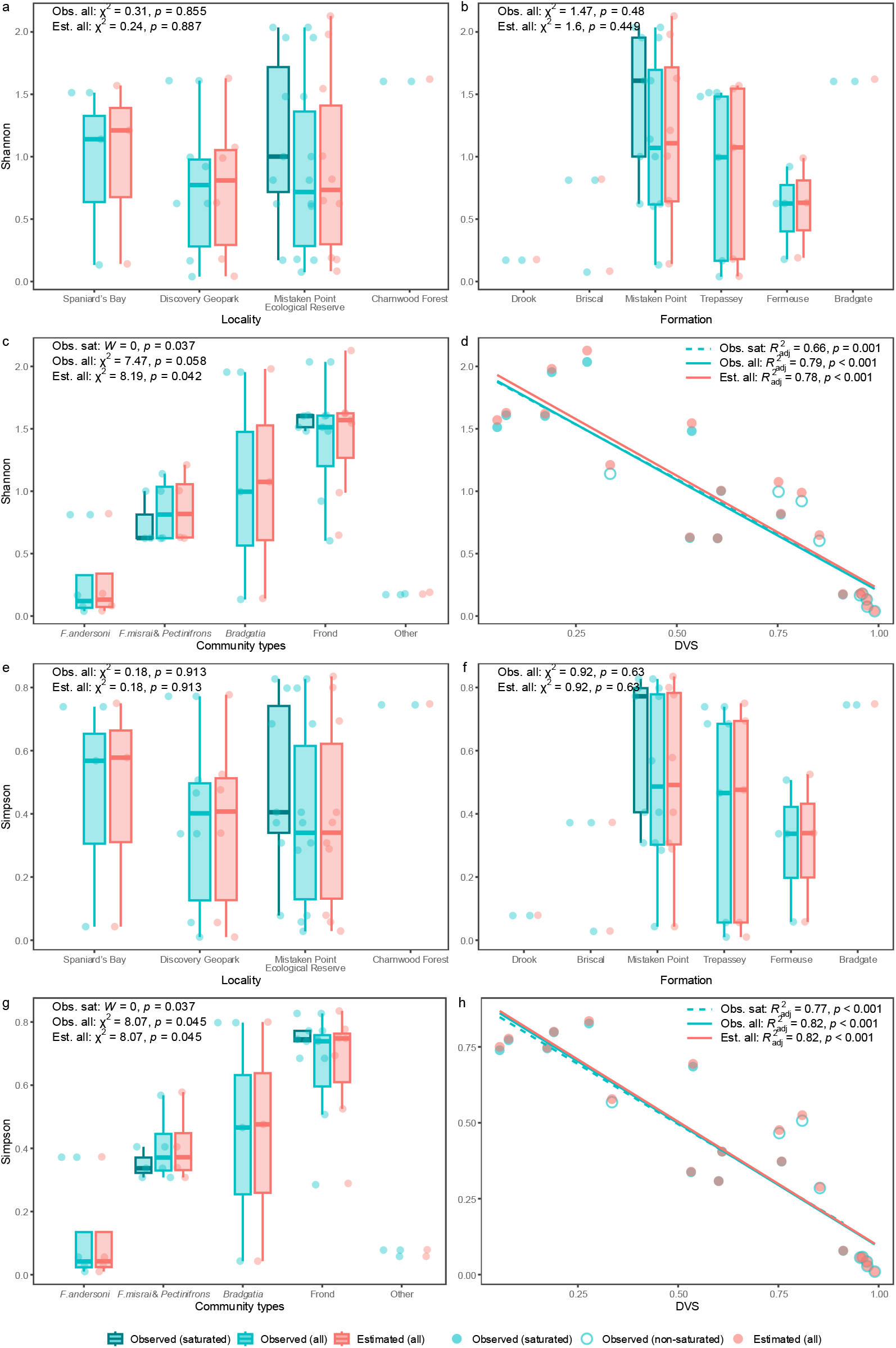
Relationships of Shannon and Simpson diversity with candidate abiotic and biotic correlates after excluding the smaller duplicated outcrops. Shannon (a–d) and Simpson (e–h) diversity were compared among (a,e) localities, (b,f) correlated formations and (c,g) community types, and analysed in relation to (d,h) community-level DVS, using observed values for saturated surfaces, observed values across all surfaces and estimated values across all surfaces. Categories represented by fewer than three surfaces were excluded from formal group comparisons. Wilcoxon rank-sum *W* and Kruskal–Wallis χ² are reported for categorical comparisons (a–c, e–g), and adjusted *R*² for linear regressions in (d,h), with corresponding *p*-values. Corresponds to Fig. 5 after excluding E Surface (Watern Cove) and G Surface (Mistaken Point).

**Figure S18.**
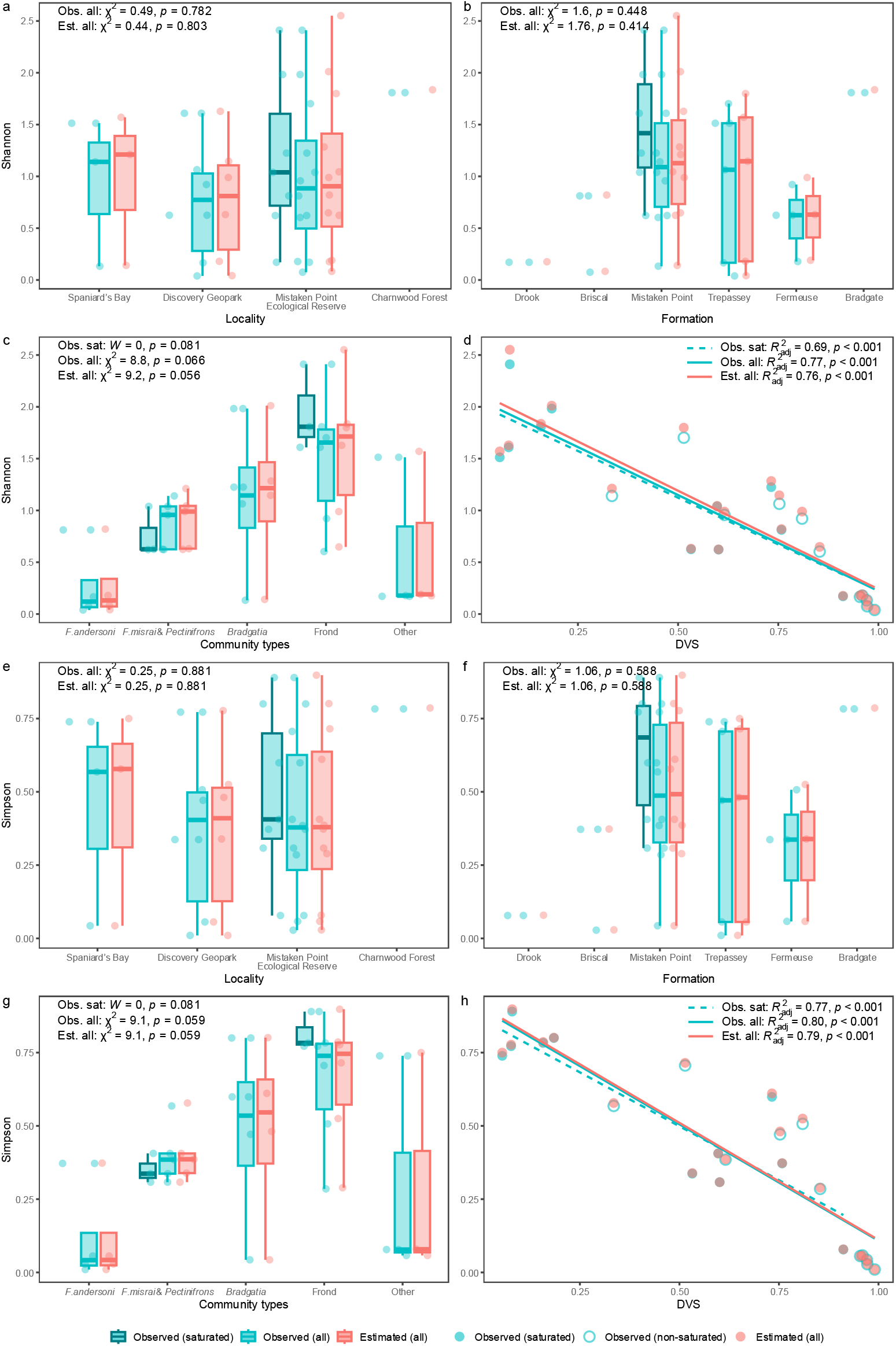
Relationships of Shannon and Simpson diversity with candidate abiotic and biotic correlates under the finer-resolution taxonomic treatment. Shannon (a–d) and Simpson (e–h) diversity were compared among (a,e) localities, (b,f) correlated formations and (c,g) community types, and analysed in relation to (d,h) community-level DVS, using observed values for saturated surfaces, observed values across all surfaces and estimated values across all surfaces. Categories represented by fewer than three surfaces were excluded from formal group comparisons. Wilcoxon rank-sum *W* and Kruskal–Wallis χ² are reported for categorical comparisons (a–c, e–g), and adjusted *R*² for linear regressions in (d,h), with corresponding *p*- values. Corresponds to Fig. 5, with specimens within the “beothukiid”, *Charniodiscus* and *Primocandelabrum* groupings resolved to species level where possible.

**Table S1.** Surface metadata and stratigraphic assignments. Locality, local stratigraphic unit and correlated formation are reported for each surface. Correlated formations follow published regional lithostratigraphic correlations ^33^.

| **Surface** | **Locality** | **Local stratigraphic unit** | **Correlated formation** |
|---|---|---|---|
| Bed B | Charnwood Forest | Bradgate Formation | Bradgate Formation |
| Bishop's Cove | Spaniard’s Bay | Mistaken Point Formation | Mistaken Point Formation |
| Brasier | Mistaken Point Ecological Reserve | Briscal Formation | Briscal Formation |
| Bristy Cove | Mistaken Point Ecological Reserve | Briscal Formation | Briscal Formation |
| Caplin Gulch | Discovery Geopark | Fermeuse Formation | Fermeuse Formation |
| Clapham's Pigeon Cove | Mistaken Point Ecological Reserve | Drook Formation | Drook Formation |
| D Surface | Mistaken Point Ecological Reserve | Mistaken Point Formation | Mistaken Point Formation |
| E Surface (Mistaken Point) | Mistaken Point Ecological Reserve | Mistaken Point Formation | Mistaken Point Formation |
| E Surface (Watern Cove) | Mistaken Point Ecological Reserve | Mistaken Point Formation | Mistaken Point Formation |
| G Surface (Mistaken Point) | Mistaken Point Ecological Reserve | Mistaken Point Formation | Mistaken Point Formation |
| G Surface (Watern Cove) | Mistaken Point Ecological Reserve | Mistaken Point Formation | Mistaken Point Formation |
| Goldmine | Discovery Geopark | Catalina Member | Trepassey Formation |
| Green Point | Spaniard’s Bay | Mistaken Point Formation | Mistaken Point Formation |
| H5 | Discovery Geopark | Murphy's Cove Member | Mistaken Point Formation |
| H14 Johnson | Discovery Geopark | Catalina Member | Trepassey Formation |
| H26 Primocandelabrum | Discovery Geopark | Fermeuse Formation | Fermeuse Formation |
| Lower Mistaken Point | Mistaken Point Ecological Reserve | Mistaken Point Formation | Mistaken Point Formation |
| Pizzeria | Mistaken Point Ecological Reserve | Trepassey Formation | Trepassey Formation |
| Shingle Head | Mistaken Point Ecological Reserve | Fermeuse Formation | Fermeuse Formation |
| St. Shotts | Mistaken Point Ecological Reserve | Mistaken Point Formation | Mistaken Point Formation |
| The Factory (H31) | Discovery Geopark | Port Union Member | Trepassey Formation |
| Upper Island Cove | Spaniard’s Bay | Trepassey Formation | Trepassey Formation |

**Table S2.** Surface sampling frame and biological summary. Original mapped area, retrodeformed mapped area, retrodeformation factor, number of specimens, specimen density and taxonomic richness are reported for each surface. Density was calculated using retrodeformed area. For taxonomic richness under the finer-resolution treatment, specimens within the “beothukiid”, *Charniodiscus* and *Primocandelabrum* groupings were resolved to species level where possible.

| **Surface** | **Mapped area (m<sup>2</sup>)** | **Retrodeformed area (m<sup>2</sup>)** | **Retrodeformation factor** | **Number of specimens (*n*)** | **Specimen density (*n* / m<sup>2</sup>)** | **Taxonomic richness (main)** | **Taxonomic richness (finer)** |
|---|---|---|---|---|---|---|---|
| Bed B | 114.56 | 114.56 | 1.00 | 272 | 2.37 | 9 | 12 |
| Bishop's Cove | 31.30 | 37.34 | 1.19 | 364 | 9.75 | 5 | 5 |
| Brasier | 40.86 | 48.57 | 1.19 | 560 | 11.53 | 8 | 8 |
| Bristy Cove | 0.96 | 0.96 | 1.00 | 70 | 73.06 | 2 | 2 |
| Caplin Gulch | 11.23 | 11.23 | 1.00 | 175 | 15.58 | 3 | 3 |
| Clapham's Pigeon Cove | 33.83 | 33.83 | 1.00 | 98 | 2.90 | 2 | 2 |
| D Surface | 72.88 | 122.48 | 1.68 | 1756 | 14.34 | 8 | 8 |
| E Surface (Mistaken Point) | 82.93 | 134.82 | 1.63 | 2314 | 17.16 | 14 | 17 |
| E Surface (Watern Cove) | 20.08 | 20.08 | 1.00 | 332 | 16.53 | 12 | 14 |
| G Surface (Mistaken Point) | 7.22 | 10.91 | 1.51 | 49 | 4.49 | 6 | 6 |
| G Surface (Watern Cove) | 93.53 | 134.68 | 1.44 | 488 | 3.62 | 18 | 20 |
| Goldmine | 14.27 | 20.97 | 1.47 | 212 | 10.11 | 5 | 5 |
| Green Point | 9.45 | 13.29 | 1.41 | 55 | 4.14 | 6 | 6 |
| H5 | 4.58 | 6.75 | 1.47 | 164 | 24.28 | 7 | 7 |
| H14 Johnson | 82.40 | 82.40 | 1.00 | 3749 | 45.50 | 9 | 9 |
| H26 Primocandelabrum | 6.96 | 9.33 | 1.34 | 29 | 3.11 | 4 | 4 |
| Lower Mistaken Point | 21.36 | 33.05 | 1.55 | 111 | 3.36 | 13 | 17 |
| Pizzeria | 70.42 | 70.42 | 1.00 | 79 | 1.12 | 8 | 11 |
| Shingle Head | 76.50 | 123.39 | 1.61 | 273 | 2.21 | 6 | 6 |
| St. Shotts | 34.36 | 55.44 | 1.61 | 74 | 1.33 | 5 | 5 |
| The Factory (H31) | 7.37 | 8.24 | 1.12 | 49 | 5.95 | 6 | 7 |
| Upper Island Cove | 16.40 | 16.40 | 1.00 | 69 | 4.21 | 7 | 7 |

**Table S3.** Sampling-completeness estimates. Observed taxonomic richness, best-fitting asymptotic model, estimated asymptotic richness and saturation classification are reported for each surface. The model with the lowest AIC was selected, and surfaces reaching at least 80% of the estimated asymptotic richness were classified as saturated. MM2/MM3 = two-/three-parameter Michaelis–Menten models; AR2/AR3 = two-/three-parameter asymptotic regression models.

| **Surface** | **Observed richness** | **Asymptote richness** | **Best model** | **Saturated** |
|---|---|---|---|---|
| Bed B | 9.00 | 9.30 | MM3 | Yes |
| Bishop's Cove | 5.00 | 6.48 | AR3 | No |
| Brasier | 8.00 | 8.66 | MM3 | Yes |
| Bristy Cove | 2.00 | 9.48 | AR3 | No |
| Caplin Gulch | 3.00 | 3.00 | AR3 | Yes |
| Clapham's Pigeon Cove | 2.00 | 2.06 | AR3 | Yes |
| D Surface | 8.00 | 9.80 | MM3 | Yes |
| E Surface (Mistaken Point) | 14.00 | 14.92 | MM3 | Yes |
| E Surface (Watern Cove) | 12.00 | 16.59 | MM3 | No |
| G Surface (Mistaken Point) | 6.00 | 7.40 | MM3 | Yes |
| G Surface (Watern Cove) | 18.00 | 20.68 | MM3 | Yes |
| Goldmine | 5.00 | 7.35 | AR3 | No |
| Green Point | 6.00 | 7.81 | MM3 | No |
| H5 | 7.00 | 7.16 | MM2 | Yes |
| H14 Johnson | 9.00 | 23.41 | MM3 | No |
| H26 Primocandelabrum | 4.00 | 5.73 | MM3 | No |
| Lower Mistaken Point | 13.00 | 15.48 | MM3 | Yes |
| Pizzeria | 8.00 | 9.47 | MM3 | Yes |
| Shingle Head | 6.00 | 9.88 | AR3 | No |
| St. Shotts | 5.00 | 7.82 | MM3 | No |
| The Factory (H31) | 6.00 | 11.02 | MM3 | No |
| Upper Island Cove | 7.00 | 8.25 | MM3 | Yes |

**Table S4.** Sampling-completeness estimates under the finer-resolution taxonomic treatment. Observed taxonomic richness, best-fitting asymptotic model, estimated asymptotic richness and saturation classification are reported for each surface. The model with the lowest AIC was selected, and surfaces reaching at least 80% of the estimated asymptotic richness were classified as saturated. Corresponds to Table S3, with specimens within the “beothukiid”, *Charniodiscus* and *Primocandelabrum* groupings resolved to species level where possible. MM2/MM3 = two-/three-parameter Michaelis–Menten models; AR2/AR3 = two-/three-parameter asymptotic regression models.

| Surface | Observed richness | Asymptote richness | Best model | Saturated |
|---|---|---|---|---|
| Bed B | 12.00 | 12.86 | MM3 | Yes |
| Bishop's Cove | 5.00 | 6.38 | AR3 | No |
| Brasier | 8.00 | 8.65 | MM3 | Yes |
| Bristy Cove | 2.00 | 8.63 | AR3 | No |
| Caplin Gulch | 3.00 | 3.00 | AR3 | Yes |
| Clapham's Pigeon Cove | 2.00 | 2.06 | AR3 | Yes |
| D Surface | 8.00 | 9.73 | MM3 | Yes |
| E Surface (Mistaken Point) | 17.00 | 18.19 | MM3 | Yes |
| E Surface (Watern Cove) | 14.00 | 20.21 | MM3 | No |
| G Surface (Mistaken Point) | 6.00 | 7.42 | MM3 | Yes |
| G Surface (Watern Cove) | 20.00 | 23.97 | MM3 | Yes |
| Goldmine | 5.00 | 7.28 | AR3 | No |
| Green Point | 6.00 | 7.77 | MM3 | No |
| H5 | 7.00 | 7.16 | MM2 | Yes |
| H14 Johnson | 9.00 | 23.54 | MM3 | No |
| H26 Primocandelabrum | 4.00 | 5.64 | MM3 | No |
| Lower Mistaken Point | 17.00 | 20.54 | MM3 | Yes |
| Pizzeria | 11.00 | 13.87 | MM3 | No |
| Shingle Head | 6.00 | 9.85 | AR3 | No |
| St. Shotts | 5.00 | 7.72 | MM3 | No |
| The Factory (H31) | 7.00 | 17.08 | MM3 | No |
| Upper Island Cove | 7.00 | 8.27 | MM3 | Yes |

**Table S5.** Estimated diversity and uncertainty for each surface.

| **Surface** | **Richness** |  |  |  | **Shannon** |  |  |  | **Simpson** |  |  |  | **Evenness** |  |  |  |
|---|---|---|---|---|---|---|---|---|---|---|---|---|---|---|---|---|
|  | Obs. | Est. | 95%<br>Lower | 95%<br>Upper | Obs. | Est. | 95%<br>Lower | 95%<br>Upper | Obs. | Est. | 95%<br>Lower | 95%<br>Upper | Obs. | Est. | 95%<br>Lower | 95%<br>Upper |
| Bed B | 9.00 | 10.00 | 9.07 | 22.91 | 1.60 | 1.62 | 1.60 | 1.72 | 0.75 | 0.75 | 0.75 | 0.78 | 0.73 | 0.70 | 0.56 | 0.85 |
| Bishop's Cove | 5.00 | 6.00 | 5.07 | 18.47 | 0.13 | 0.14 | 0.13 | 0.22 | 0.04 | 0.04 | 0.04 | 0.07 | 0.08 | 0.08 | 0.03 | 0.13 |
| Brasier | 8.00 | 9.00 | 8.07 | 21.86 | 0.81 | 0.82 | 0.81 | 0.92 | 0.37 | 0.37 | 0.37 | 0.42 | 0.39 | 0.37 | 0.28 | 0.47 |
| Bristy Cove | 2.00 | 2.00 | 2.00 | 3.60 | 0.08 | 0.08 | 0.08 | 0.21 | 0.03 | 0.03 | 0.03 | 0.08 | 0.11 | 0.12 | 0.00 | 0.32 |
| Caplin Gulch | 3.00 | 3.00 | 3.00 | 3.01 | 0.63 | 0.63 | 0.63 | 0.75 | 0.34 | 0.34 | 0.34 | 0.42 | 0.57 | 0.57 | 0.47 | 0.68 |
| Clapham's Pigeon Cove | 2.00 | 2.00 | 2.00 | 2.28 | 0.17 | 0.18 | 0.17 | 0.31 | 0.08 | 0.08 | 0.08 | 0.15 | 0.25 | 0.25 | 0.06 | 0.45 |
| D Surface | 8.00 | 10.00 | 8.18 | 30.12 | 0.62 | 0.62 | 0.62 | 0.67 | 0.31 | 0.31 | 0.31 | 0.33 | 0.30 | 0.27 | 0.18 | 0.36 |
| E Surface (Mistaken Point) | 14.00 | 17.00 | 14.35 | 39.42 | 1.00 | 1.01 | 1.00 | 1.06 | 0.41 | 0.41 | 0.41 | 0.43 | 0.38 | 0.35 | 0.29 | 0.42 |
| E Surface (Watern Cove) | 12.00 | 15.99 | 12.56 | 40.64 | 0.92 | 0.95 | 0.92 | 1.09 | 0.38 | 0.39 | 0.38 | 0.45 | 0.37 | 0.34 | 0.25 | 0.44 |
| G Surface (Mistaken Point) | 6.00 | 6.25 | 6.01 | 10.66 | 1.23 | 1.28 | 1.23 | 1.57 | 0.60 | 0.61 | 0.60 | 0.74 | 0.68 | 0.70 | 0.52 | 0.88 |
| G Surface (Watern Cove) | 18.00 | 25.98 | 18.99 | 82.69 | 1.95 | 1.98 | 1.95 | 2.08 | 0.80 | 0.80 | 0.80 | 0.82 | 0.68 | 0.61 | 0.44 | 0.77 |
| Goldmine | 5.00 | 6.00 | 5.09 | 16.02 | 0.17 | 0.18 | 0.17 | 0.29 | 0.06 | 0.06 | 0.06 | 0.10 | 0.10 | 0.10 | 0.03 | 0.17 |
| Green Point | 6.00 | 7.96 | 6.18 | 27.76 | 1.14 | 1.21 | 1.14 | 1.53 | 0.57 | 0.58 | 0.57 | 0.70 | 0.64 | 0.58 | 0.29 | 0.88 |
| H5 | 7.00 | 7.00 | 7.00 | 8.39 | 1.61 | 1.63 | 1.61 | 1.73 | 0.77 | 0.78 | 0.77 | 0.80 | 0.83 | 0.84 | 0.76 | 0.91 |
| H14 Johnson | 9.00 | 17.00 | 9.99 | 73.80 | 0.04 | 0.04 | 0.04 | 0.06 | 0.01 | 0.01 | 0.01 | 0.01 | 0.02 | 0.01 | 0.01 | 0.02 |
| H26 Primocandelabrum | 4.00 | 4.48 | 4.03 | 12.20 | 0.92 | 0.99 | 0.92 | 1.31 | 0.51 | 0.53 | 0.51 | 0.71 | 0.66 | 0.66 | 0.33 | 0.99 |
| Lower Mistaken Point | 13.00 | 20.93 | 13.98 | 77.26 | 2.04 | 2.13 | 2.04 | 2.32 | 0.83 | 0.84 | 0.83 | 0.88 | 0.79 | 0.70 | 0.44 | 0.96 |
| Pizzeria | 8.00 | 9.98 | 8.18 | 29.87 | 1.48 | 1.55 | 1.48 | 1.76 | 0.69 | 0.69 | 0.69 | 0.78 | 0.71 | 0.67 | 0.44 | 0.90 |
| Shingle Head | 6.00 | 6.66 | 6.06 | 13.78 | 0.18 | 0.19 | 0.18 | 0.31 | 0.06 | 0.06 | 0.06 | 0.10 | 0.10 | 0.10 | 0.03 | 0.17 |
| St. Shotts | 5.00 | 6.97 | 5.18 | 26.86 | 0.60 | 0.65 | 0.60 | 0.92 | 0.29 | 0.29 | 0.29 | 0.41 | 0.37 | 0.33 | 0.11 | 0.56 |
| The Factory (H31) | 6.00 | 7.96 | 6.18 | 27.72 | 1.00 | 1.08 | 1.00 | 1.43 | 0.47 | 0.48 | 0.47 | 0.64 | 0.56 | 0.52 | 0.23 | 0.80 |
| Upper Island Cove | 7.00 | 7.99 | 7.07 | 20.61 | 1.51 | 1.57 | 1.51 | 1.74 | 0.74 | 0.75 | 0.74 | 0.80 | 0.78 | 0.76 | 0.54 | 0.97 |

**Table S6.**
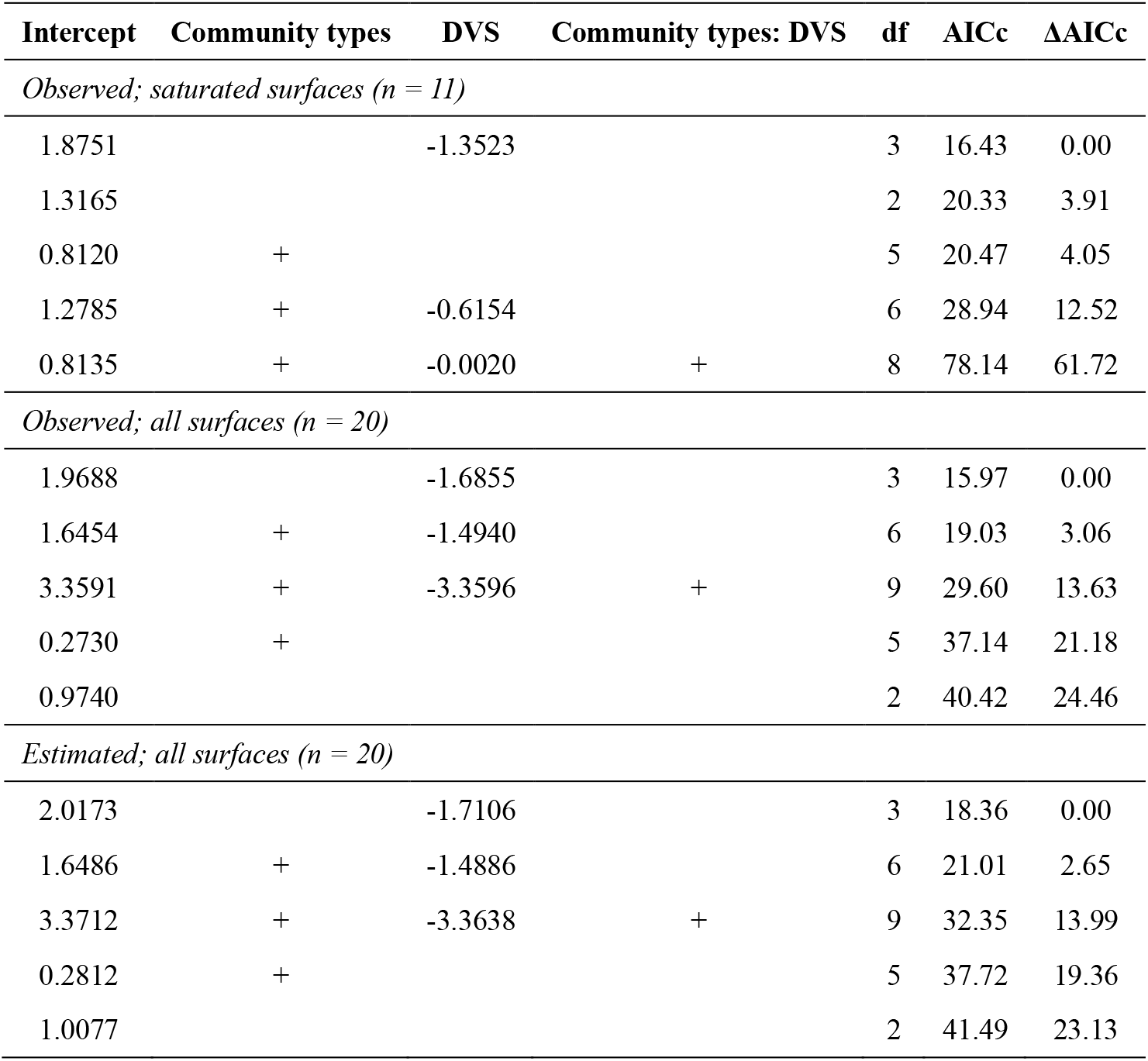
AICc comparison of models relating Shannon diversity to community type and DVS. Candidate models were derived from the global model *Shannon diversity ∼ community type × DVS* and fitted separately to observed diversity among saturated surfaces, observed diversity across all surfaces and estimated diversity across all surfaces. Analyses included the four principal community types only. Lower AICc values indicate greater model support.

## References

1. Stein, A., Gerstner, K. & Kreft, H. Environmental heterogeneity as a universal driver of species richness across taxa, biomes and spatial scales. Ecology Letters 17, 866–880 (2014).

2. Leibold, M. A. et al. The metacommunity concept: a framework for multi-scale community ecology. Ecology Letters 7, 601–613 (2004).

3. Tilman, D. Competition and biodiversity in spatially structured habitats. Ecology 75, 2–16 (1994).

4. Cornell, H. V. & Harrison, S. P. What are species pools and when are they important? Annual Review of Ecology, Evolution, and Systematics vol. 45 45–67 (2014).

5. Mittelbach, G. G. & Schemske, D. W. Ecological and evolutionary perspectives on community assembly. Trends in Ecology & Evolution 30, 241–247 (2015).

6. Vellend, M. Conceptual synthesis in community ecology. The Quarterly Review of Biology 85, 183–206 (2010).

7. HilleRisLambers, J., Adler, P. B., Harpole, W. S., Levine, J. M. & Mayfield, M. M. Rethinking community assembly through the lens of coexistence theory. Annual Review of Ecology, Evolution, and Systematics vol. 43 227–248 (2012).

8. Bruno, J. F., Stachowicz, J. J. & Bertness, M. D. Inclusion of facilitation into ecological theory. Trends in Ecology & Evolution 18, 119–125 (2003).

9. Bregman, T. P. et al. Species interactions regulate the collapse of biodiversity and ecosystem function in tropical forest fragments. Ecology 96, 2692–2704 (2015).

10. Butterfield, N. J. Macroevolution and macroecology through deep time. Palaeontology 50, 41–55 (2007).

11. Darroch, S. A. F., Laflamme, M. & Wagner, P. J. High ecological complexity in benthic Ediacaran communities. Nature Ecology & Evolution 2, 1541–1547 (2018).

12. Liu, A. G., Kenchington, C. G. & Mitchell, E. G. Remarkable insights into the paleoecology of the Avalonian Ediacaran macrobiota. Gondwana Research 27, 1355–1380 (2015).

13. Mitchell, E. G. & Pates, S. From organisms to biodiversity: The ecology of the Ediacaran/Cambrian transition. Paleobiology 51, 150–173 (2025).

14. Benus, A., Landing, E., Narbonne, G. & Myrow, P. Sedimentological context of a deep-water Ediacaran fauna (Mistaken Point, Avalon Zone, eastern Newfoundland). Trace fossils, small shelly fossils and the Precambrian–Cambrian boundary 463, 8–9 (1988).

15. Wood, D. A., Dalrymple, R. W., Narbonne, G. M., Gehling, J. G. & Clapham, M. E. Paleoenvironmental analysis of the late Neoproterozoic Mistaken Point and Trepassey formations, southeastern Newfoundland. Canadian Journal of Earth Sciences 40, 1375–1391 (2003).

16. Clapham, M. E., Narbonne, G. M. & Gehling, J. G. Paleoecology of the oldest known animal communities: Ediacaran assemblages at Mistaken Point, Newfoundland. Paleobiology 29, 527–544 (2003).

17. Stephenson, N. P. et al. Morphology shapes community dynamics in early animal ecosystems. Nature Ecology & Evolution 8, 1238–1247 (2024).

18. Finnegan, S., Gehling, J. G. & Droser, M. L. Unusually variable paleocommunity composition in the oldest metazoan fossil assemblages. Paleobiology 45, 235–245 (2019).

19. Eden, R., Manica, A. & Mitchell, E. G. Metacommunity analyses show an increase in ecological specialisation throughout the Ediacaran period. PLOS Biology 20, e3001289 (2022).

20. Stephenson, N. P. et al. Recolonization strategies of early animals in the Avalon (Ediacaran 574–560 Ma). Paleobiology 52, 57–68 (2026).

21. Clapham, M. E. & Narbonne, G. M. Ediacaran epifaunal tiering. Geology 30, 627–630 (2002).

22. Mitchell, E. G. & Kenchington, C. G. The utility of height for the Ediacaran organisms of Mistaken Point. Nature Ecology & Evolution 2, 1218–1222 (2018).

23. Mitchell, E. G. et al. The importance of neutral over niche processes in structuring Ediacaran early animal communities. Ecology Letters 22, 2028–2038 (2019).

24. Mitchell, E. G. & Manica, A. The influence of reproductive mode on resource competition and diversity patterns in Ediacaran early animal communities. Nature Ecology & Evolution https://doi.org/10.1038/s41559-026-03094-2 (2026) doi:10.1038/s41559-026-03094-2.

25. Matthews, J. J. et al. A chronostratigraphic framework for the rise of the Ediacaran macrobiota: New constraints from Mistaken Point Ecological Reserve, Newfoundland. GSA Bulletin 133, 612–624 (2021).

26. Bottjer, D. J. & Ausich, W. I. Phanerozoic development of tiering in soft substrata suspension-feeding communities. Paleobiology 12, 400–420 (1986).

27. Chao, A. et al. Rarefaction and extrapolation with Hill numbers: a framework for sampling and estimation in species diversity studies. Ecological Monographs 84, 45–67 (2014).

28. Gotelli, N. J. & Colwell, R. K. Quantifying biodiversity: procedures and pitfalls in the measurement and comparison of species richness. Ecology Letters 4, 379–391 (2001).

29. Barnosky, A. D., Carrasco, M. A. & Davis, E. B. The impact of the species–area relationship on estimates of paleodiversity. PLOS Biology 3, e266 (2005).

30. Colwell, R. K. et al. Models and estimators linking individual-based and sample-based rarefaction, extrapolation and comparison of assemblages. Journal of Plant Ecology 5, 3–21 (2012).

31. Hofmann, H. J., O’Brien, S. J. & King, A. F. Ediacaran biota on Bonavista Peninsula, Newfoundland, Canada. Journal of Paleontology 82, 1–36 (2008).

32. Ichaso, A. A., Dalrymple, R. W. & Narbonne, G. M. Paleoenvironmental and basin analysis of the late Neoproterozoic (Ediacaran) upper Conception and St. John’s groups, west Conception Bay, Newfoundland. Can. J. Earth Sci. 44, 25–41 (2007).

33. Mason, S. J., Narbonne, G. M., Dalrymple, R. W. & O’Brien, S. J. Paleoenvironmental analysis of Ediacaran strata in the Catalina Dome, Bonavista Peninsula, Newfoundland. Can. J. Earth Sci. 50, 197–212 (2013).

34. King, A. F. Geology of the Avalon Peninsula, Newfoundland (parts of 1K, 1L, 1M, 1N and 2C). (1988).

35. Mitchell, E. G. et al. Variation of population and community ecology over large spatial scales in Ediacaran early animal communities. Global and Planetary Change 251, 104818 (2025).

36. Kenchington, C. G., Harris, S. J., Vixseboxse, P. B., Pickup, C. & Wilby, P. R. The Ediacaran fossils of Charnwood Forest: Shining new light on a major biological revolution. Proceedings of the Geologists’ Association 129, 264–277 (2018).

37. Rosse-Guillevic, S., Muirhead-Hunt, H. & McIlroy, D. Aninoides: a new rangeomorph genus from the upper Ediacaran of Newfoundland. Papers in Palaeontology 11, e70061 (2025).

38. Boynton, H. & Ford, T. Ediacaran fossils from the precambrian (charnian supergroup) of Charnwood Forest, Leicestershire, England. Mercian Geologist 13, 165–182 (1995).

39. Oksanen, J., et al. Vegan: Community Ecology Package. (2026). doi:10.32614/CRAN.package.vegan.

40. Suzuki, R., Terada, Y. & Shimodaira, H. Pvclust: Hierarchical Clustering with p-Values via Multiscale Bootstrap Resampling. (2019). doi:10.32614/CRAN.package.pvclust.

41. Rousseeuw, P. J. Silhouettes: A graphical aid to the interpretation and validation of cluster analysis. Journal of Computational and Applied Mathematics 20, 53–65 (1987).

42. Maechler, M., Rousseeuw, P., Struyf, A., Hubert, M. & Hornik, K. Cluster: Cluster Analysis Basics and Extensions. (2026).

43. Laflamme, M., Xiao, S. & Kowalewski, M. Osmotrophy in modular Ediacara organisms. Proceedings of the National Academy of Sciences 106, 14438–14443 (2009).

44. Laflamme, M., Flude, L. I. & Narbonne, G. M. Ecological tiering and the evolution of a stem: The oldest stemmed frond from the Ediacaran of Newfoundland, Canada. Journal of Paleontology 86, 193–200 (2012).

45. Narbonne, G. M. Modular construction of early Ediacaran complex life forms. Science 305, 1141–1144 (2004).

46. Baddeley, A. & Turner, R. spatstat: An R package for analyzing spatial point patterns. J. Stat. Soft. 12, 1–42 (2005).

47. Baddeley, A., Rubak, E. & Turner, R. Spatial Point Patterns: Methodology and Applications with R. (CRC press Boca Raton, 2015).

48. R Core Team. R: A Language and Environment for Statistical Computing. (R Foundation for Statistical Computing, Vienna, Austria, 2026). doi:10.32614/R.manuals.

49. Ritz, C., Baty, F., Streibig, J. C. & Gerhard, D. Dose-response analysis using R. PLOS ONE 10, e0146021 (2016).

50. Hsieh, T. C., Ma, K. H. & Chao, A. iNEXT: Interpolation and Extrapolation for Species Diversity. (2025).

51. Oehlert, G. W. A note on the delta method. The American Statistician 46, 27–29 (1992).

52. Bartoń, K. MuMIn: Multi-Model Inference. (2026). doi:10.32614/CRAN.package.MuMIn.

53. Lin, L. I.-K. A concordance correlation coefficient to evaluate reproducibility. Biometrics 45, 255–268 (1989).

54. Signorell, A. DescTools: Tools for Descriptive Statistics. (2025). doi:10.32614/CRAN.package.DescTools.

55. McBride, G. B. A Proposal for Strength-of-Agreement Criteria for Lin’s Concordance Correlation Coefficient. (2005).

56. DeMalach, N., Saiz, H., Zaady, E. & Maestre, F. T. Plant species–area relationships are determined by evenness, cover and aggregation in drylands worldwide. Global Ecology and Biogeography 28, 290–299 (2019).

57. Daly, A. J., Baetens, J. M. & De Baets, B. Ecological diversity: Measuring the unmeasurable. Mathematics 6, 119 (2018).

58. Roswell, M., Dushoff, J. & Winfree, R. A conceptual guide to measuring species diversity. Oikos 130, 321–338 (2021).

59. Chesson, P. Mechanisms of maintenance of species diversity. Annual Review of Ecology, Evolution, and Systematics vol. 31 343–366 (2000).

60. Adler, P. B., HilleRisLambers, J. & Levine, J. M. A niche for neutrality. Ecology Letters 10, 95–104 (2007).

